# Planning in Nonhuman Primates Emerges from Structure Knowledge and is Distinct from Attention, Working Memory, Effort Control, and Learning

**DOI:** 10.64898/2026.07.17.739240

**Authors:** Xuan Wen, Adam Neumann, Seema Dhungana, Paul Tiesinga, Thilo Womelsdorf

## Abstract

Forward planning involves mentally simulating future choices. Such mental simulation may constitute an independent cognitive operation, or it may be intricately linked to canonical cognitive operations supporting working memory of choice options, covert attentional information sampling, forming memory models of problem structure and effort control for pursuing future goals. Here, we test the relative independence of planning in non-human primates with an object-sequence planning task assessed together with an assay of simpler cognitive tasks. The sequence task unexpectedly masked objects at future serial positions probing NHP’s to plan ahead serial future choices. We found that five of six subjects planned ahead 2-3 ordinal positions. Planning depth was predicted by knowledge of the serial structure but constituted a unique cognitive factor that was distinct from sequence memory, object memory, and learning efficacy. Similarly, planning was segregated from working memory capacity, attention, inhibitory control and effort control. These findings suggest that planning ahead constitutes a separate construct not reducible to other domain-general cognitive operations, and emerges in NHP’s that successfully form an internal model of the problem structure.

## Introduction

The ability to plan ahead endows subjects with flexibility to reach goals that require more than one step ahead of the current behavioral state. Planning can thereby be defined as the selection of a sequence of choices to reach a favorable outcome (Mattar and Lengyel, 2022). While this definition of planning suggests that specialized mental search and inference algorithms exist that implement efficient planning (Botvinick and Toussaint, 2012; Mattar and Daw, 2026), basic planning processes involve a combination of simpler canonical cognitive operations. The simpler processes that may underlie the ability to plan includes (*i*) the ability to form an internal model of possible future choices, (*ii*) the use working memory to activate and maintain versatile future options, (*iii*) the covert sampling of information that prioritizes, i.e. attends, relevant options while suppressing non-favored options, and (*iv*) the subjective willingness to exert effort and spent cognitive resources to mentally consider future options until a versatile future option is identified. Combining these simpler canonical processes may naturally reflect mental planning operations. According this view planning may have evolved from tying together evolutionary processes that favor improvements in structure learning, working memory, attentional control and effort control (Scarf et al., 2011). Despite this possibility, it has remained unresolved how separable the abilities in these more basic cognitive processes are from the ability to plan ahead.

Studies aiming to identify the core set of cognitive factors that explain performance variance on executive control tasks in humans have not found planning to be a distinct factor. Rather, planning abilities are subsumed within a common cognitive control factor, or spread across more basic cognitive factors that include working memory updating, shifting and inhibitory control processes (e.g. (Engle et al., 1999; Miyake et al., 2000)). Similarly, when planning has been studied in monkeys, the limitations of planning abilities are rarely linked to constraints in planning abilities per se. Rather, limitations to plan ahead not much more than about two items in nonhuman primates are considered to reflect limitations in other cognitive operations such as limitation in working memory (WM) capacity that constrain the number of future items that can be maintained online (Scarf et al., 2011; Templer et al., 2019; Wen et al., 2025b); limitation in attentional control and inhibition that constrain the ability to suppress sub-goals that are non-favorable and that would increase the distance from a desired future goal (Fragaszy et al., 2003; Beran et al., 2015); and to overall lower levels of cognitive effort control that is needed to spent cognitive resources on considering future steps instead of directly acting out (Mattar et al., 2025). These considerations are plausible but raise an unanswered question whether planning abilities are separable at the cognitive behavioral level from abilities in WM capacity, attentional control, inhibitory control, and effort control.

Here, we address this question by testing how six nonhuman primates learn object sequences and, once learned, continue performing the sequence when objects are masked. Masks were introduced in 90% of trials after the first, second, or third correct choice of the sequence to engage subjects to spontaneously engage in forward planning to maximize the likelihood to complete the sequence and receive reward. We found that five of six subjects planned ahead by at least two objects after mask onset. The planning depth was predictable from the level of knowledge of the sequence structure, but was not associated with how fast subjects learned the sequence, how well they recalled individual objects of the sequence, or by their working memory capacity, attentional control, inhibitory control and effort control as measured with separate tasks. The results show that NHP’s spontaneously engage in planning and suggest that planning is a distinct cognitive skill that is not reducible to other cognitive factors.

## Results

Six rhesus macaques performed a sequence learning and planning task across 45-47 sessions (subject B: 45; F: 46; J: 47; K: 45; R: 45; S: 46) in which they learned to choose four objects in a pre-determined, fixed order, ignoring a fifth distracting object (**Fig. 1A**). Subjects had 20 attempts, or trials, to complete a sequence. When a sequence was completed with ≤6 errors per trial fluid reward was provided. Otherwise, the trial terminated and a new trial started with the same objects at random locations. For each new sequence the first five trials showed objects unmasked. Trials 6–20 introduced grey square masks in 90% of trials. The masks overlayed all remaining objects after the first, second, or third correct ordinal choice (**Fig. 1B**). Each session tested five sequences, followed by other tasks that tested effort control and abilities in inhibitory control and attention (**Fig. 1C**).

**Fig. 1.**
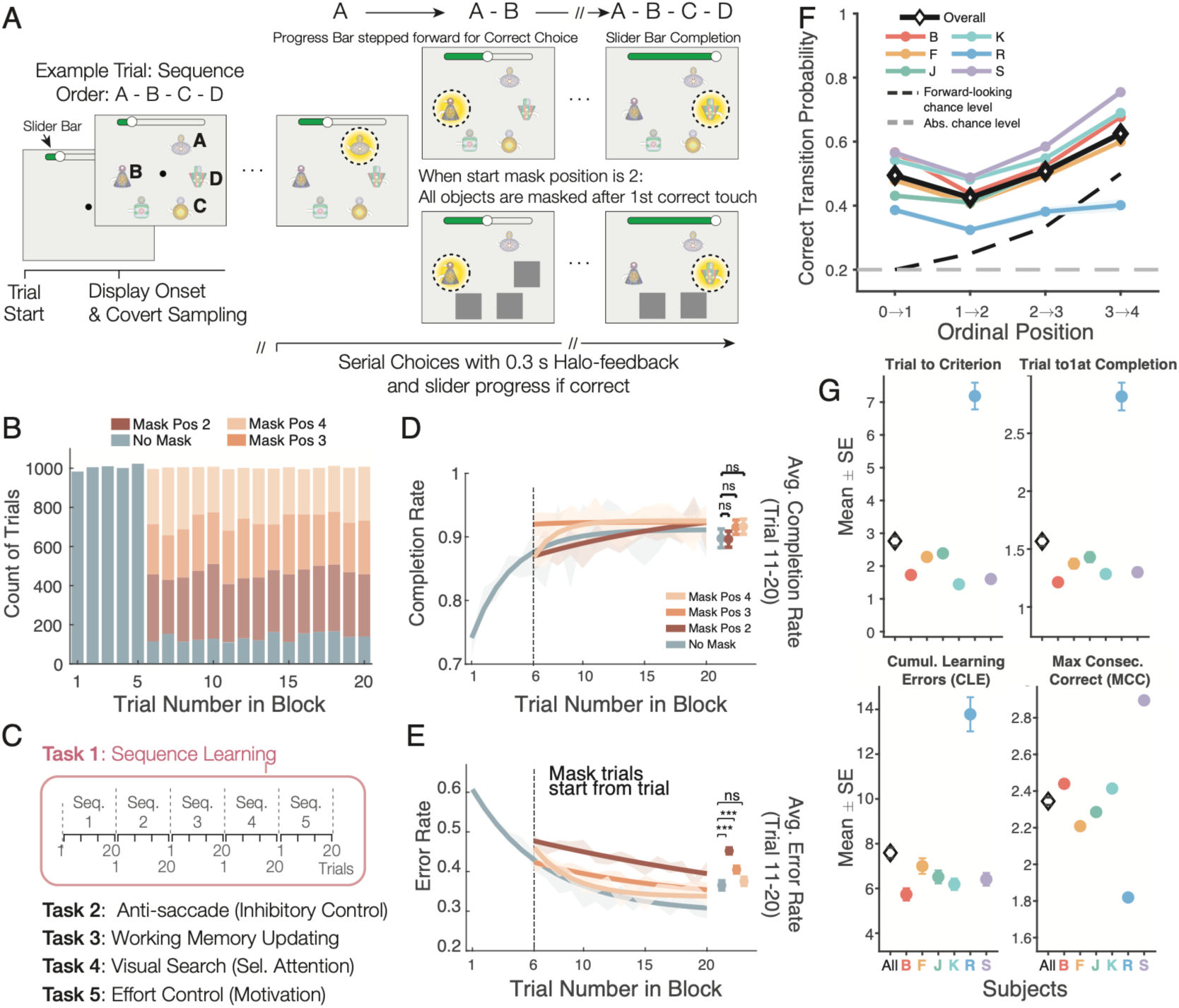
Monkeys learn and complete masked object-sequences. (**A**) Subjects chose four objects in a predefined order A-B-C-D, ignoring a fifth distractor object. A slider tracked progress through the sequence. In unmasked trials, all objects were visible; in masked trials, objects at and beyond a designated ordinal position were hidden from view. (**B**) Distribution of masking trials. Trials 1–5 were unmasked. From trial 6 onward, masks started either after the 1^st^, 2^nd^, or 3^rd^ choice, 10% trials remained unmasked. (**C**) Each session used 5 new sequences with 20 trials each. Subjects also performed antisaccade, working memory updating, visual search, and effort control tasks. (**D**) Completion rate across trials for each masking condition, fitted with exp. growth curves. Rightmost points: mean ± SE completion rate for trials 11–20 (no-mask: 0.90 ± 0.01; mask 2: 0.91 ± 0.01; mask 3: 0.92 ± 0.01; mask 4: 0.92 ± 0.01). Mask and no-mask conditions were similar (LME, all p ≥ .045; Bonf. corrected). (**E**) Same as *D* for error rates. Mask position 2 (0.44 ± 0.006, p < .001) and mask position 3 (0.40 ± 0.008, p < .001) showed higher error rates than no-mask trials (0.36 ± 0.01). (**F**) State transition probabilities at each ordinal position for no-mask trials 11–20, for each subject. Thick dashed line indicates forward-looking chance level (1 / number of remaining objects). Transition probabilities were all above both abs. chance (0.20) and forward-looking chance (state 0→1: P = 0.49, p < .001; state 1→2: P = 0.42, p < .001; state 2→3: P = 0.51, p < .001; state 3→4: P = 0.62, p = .007). (**G**) Mean (± SE) across subjects and sessions of the first trial to completion (1.57 ± 0.04), trial to criterion (2.76 ± 0.14), cumulative learning errors (7.59 ± 0.24), and max. consecutive correct trials (2.28 ± 0.02). Metrics were computed per sequence and averaged across 5 sequences of a session. Black diamond is avg. across subjects (colors).

Subjects learned the sequences, completing the first sequence on average in 1.57 (SE = 0.04) trials and continuing to reduce the number of errors in subsequent trials (**Fig. 1D,E**). From trial 5 onwards objects were masked once subjects correctly chose the first, second, or third ordinal position, which required subjects to complete the sequence without visible objects. Despite the masks, subjects continued completing sequences on trials 11–20 near ceiling across all conditions (completion rates for no-mask: 0.90 ± 0.01; masked at position 2, i.e. mask-2: 0.91 ± 0.01; mask-3: 0.92 ± 0.01; mask-4 : 0.92 ± 0.01) (**Fig. 1D**). Example performances are shown for each mask condition in **Movie S1**. However, masks overall reduced performance. Mask onsets at position 2 and 3 increased erroneous choices compared to no-mask trials (error rate without mask: 0.36; with mask-2 : 0.44 ± 0.006, p < .001; mask-3: 0.40 ± 0.008, p < .001; mask-4: 0.36 ± 0.009, n.s.; **Fig. 1E**). These results indicate that masks after the first or second choice introduced uncertainty but still allowed subjects to complete sequences without visible objects.

We next tested how masking affected choices. Without a mask, correct sequential choices in trials 11-20, measured as correct transition probabilities, were above chance at all four ordinal positions in five of six subjects and on average across subjects (state 0→1: P = 0.49, p < .001; state 1→2: P = 0.42, p < .001; state 2→3: P = 0.51, p < .001; state 3→4: P = 0.62, p = .007; **Fig. 1F**). Two-thirds of erroneously chosen objects during masking were at their wrong ordinal position. Erroneously chosen objects were more likely ‘forward’, i.e. from ordinal positions that were not-yet reached, evident in a positive ordinal signed distance of erroneously chosen objects on mask-2 (i.e. mask introduced after the first choice at position 2) and mask-3 trials (no-mask: 1.17, mask-2: 1.06, mask-3: 1.07, mask-4: 0.97; all p < .001; **Fig. S1**). This finding suggests subjects searched through the set of not-yet chosen objects prior to making a choice. The remaining errors were composed of rule-breaking errors (0.02%, e.g. failing to re-chose the last correct object after an error) and erroneous choices of the distractor object that accompanied a sequence but was never relevant (**Fig. S1C,D**). Distractor errors were faster than sequence errors (**Fig. S1E**), suggesting that distractor choices were more exploratory or impulsive than deliberative.

### Subjects plan 2-3 items ahead during masked trials

We next asked whether subjects planned ahead and correctly chose masked objects (**Fig. 2A**). If subjects planned future choices of objects before they were masked, accuracy at masked positions should exceed chance levels among the remaining objects (forward-looking chance). We aligned all masked trials by the position at which masking began and computed the difference between actual accuracy and forward-looking chance at each relative position (**Fig. 2B**). Accuracy was above chance at the first masked position (relative position 1; mean Δ = 0.21 ± 0.008, p < .001), at the 2^nd^ masked position (relative position 2; Δ = 0.045 ± 0.006, p = .006), and at the 3^rd^ masked position (relative position 3; Δ = 0.020 ± 0.008, p = .008). The steep decline in performance with the 2^nd^ and 3^rd^ masked positions could reflect the decay of passive temporal short-term memory of objects. We tested this by restricting the analysis to trials matched for retention delay across masking conditions (delay window: 0.66–3.08 s) and found that it retained the above-chance accuracy at the 2^nd^ and 3^rd^ masked position (2^nd^-masked choice: Δ = 0.032, p < .001; 3^rd^-masked choice: Δ = 0.019, p = .024). While temporal decay may still have affected the choice accuracy, it did not abolish forward planned object information within this 0.66–3.08 s window.

**Fig. 2.**
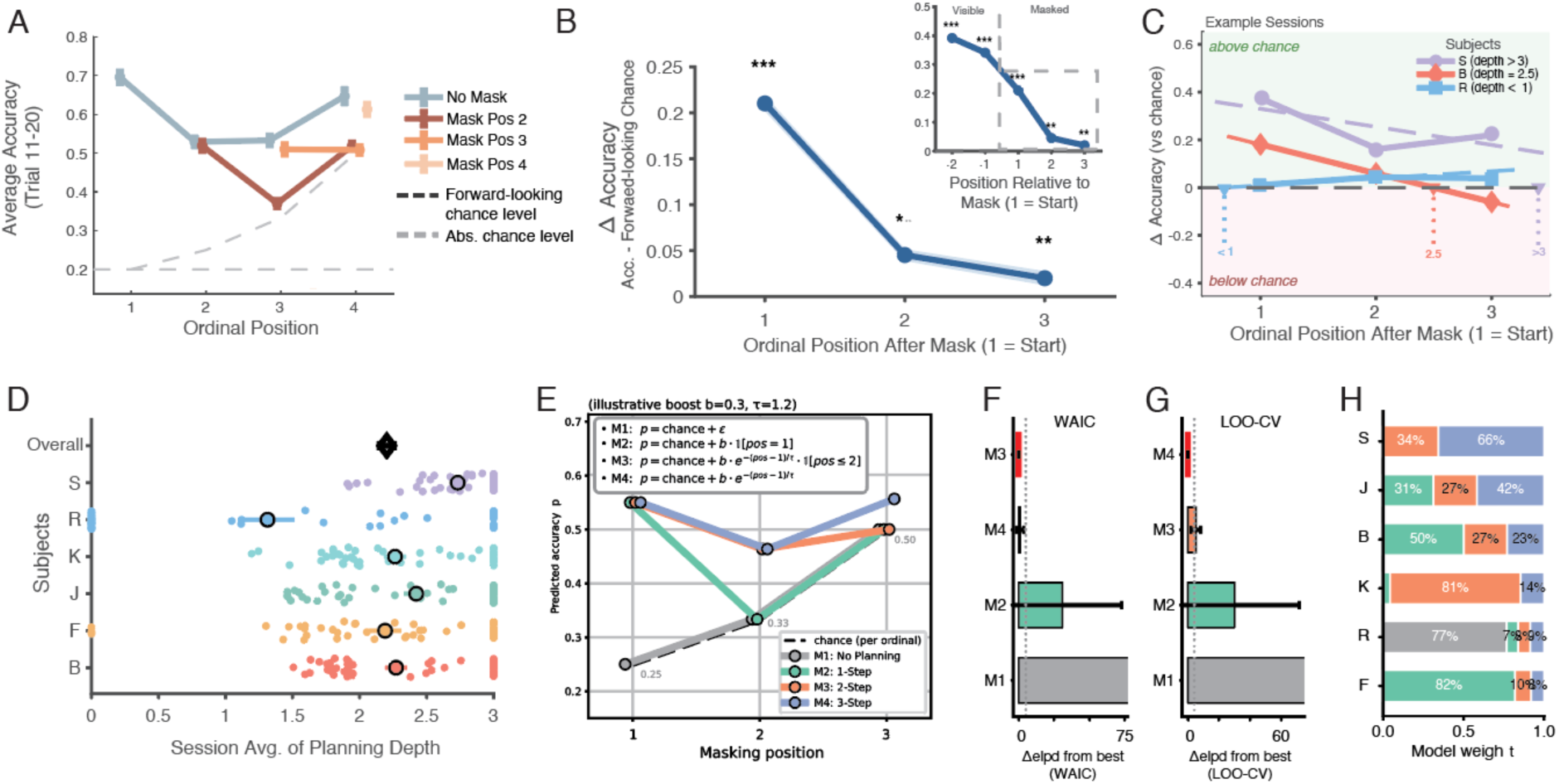
Subjects show forward planning during masked trials. (**A**) Accuracies at ordinal positions for masking conditions (trials 11–20). Dashed lines indicate chance levels. (**B**) Accuracy difference relative to forward-looking chance, aligned by mask onset position (1=1^st^ masked position). Accuracy remained above chance at 1^st^, 2^nd^ and 3^rd^ masked choice (1^st^ masked choice: mean Δ = 0.21 ± 0.008, p < .001; 2^nd^ masked choice: Δ = 0.045 ± 0.006, p = .006; 3^rd^ masked choice: Δ = 0.020 ± 0.008, p = .008). (**C**) Illustration of estimating planning depth. For each session, a three-point accuracy-minus-chance profile at relative positions 0, +1, and +2 from mask onset was linearly interpolated to find the zero-crossing, yielding a continuous planning depth estimate clipped to [0, 3]. Colored lines are example sessions from diff. subjects. (**D**) Mean (± SE) planning depth (*x-axis*) for each subject (*y-axis*). Planning depth ranged from 1.31–2.73 (avg.: = 2.20 items ahead, SE = 0.05). (**E**) Schematic of four candidate planning models: M1, no planning (chance at all masked positions); M2, one-step planning (boost only at the first masked position); M3, two-step planning (exponential decay over first two masked positions); M4, three-step planning (exponential decay over three masked positions). (**F**) WAIC (Widely Applicable Information Criterion for Bayesian model comparison) for models M1–M4. Models M3 (2-step) and M4 (3-step) provided the best fit (elpd waic = −339.4 and −340.3, respectively), outperforming M2 (1-step; Δelpd = 30.8) and M1 (no planning; Δelpd = 934.3). (**G**) Pareto-smoothed Leave-One-Out Cross-Validation (LOO-CV) for models M1–M4. Results matched those of the WAIC rankings: M4 (elpd_loo = −334.2) and M3 (−339.3) were similar, both better than M2 (−364.1) and M1 (−1219.9). (**H**) Subject-wise best-model assignment from Bayesian posterior model weights. Best-fitting models varied between subjects.

To estimate the depth of forward planning, we fitted a linear regression to the accuracy-minus-chance profile across the 1^st^, 2^nd^, and 3^rd^ ordinal choice after mask onset across sequences of individual sessions. Crossing of the chance level (i.e. zero-crossings of the fit) was then used as the estimate for the number of above-chance choices of masked objects which corresponds to planning depth as shown for examples in **Fig. 2C**. Planning depth varied across subjects, with a group mean of 2.20 (SE = 0.05) items ahead (**Fig. 2D**). At the individual level, five of six subjects showed mean planning depths between 2.19 and 2.73 items (subjects B: 2.27 ± 0.08; F: 2.19 ± 0.12; J: 2.42 ± 0.09; K: 2.26 ± 0.08; S: 2.73 ± 0.05), while one subject (R) showed markedly lower depth of 1.31 ± 0.20 objects after mask onset. We used leave-one-subject-out analyses to validate that no single subject drove the group-level effect with all six leave-one-out group means remaining between 2.09 and 2.37 (**Fig. S1J-K**).

We next used a model-based approach to retrieve the planning strategies of individual subjects by fitting Bayesian models to the data of the masked trials of each subject: A first model M1 assumed no planning, predicting accuracy at chance for all masked positions; model M2 assumed a one-step look-ahead, predicting above-chance accuracy only at the first masked position (green line in **Fig. 2E**); M3 assumed a two-step exponential decay in retrieval probability across masked positions; and M4 assumed a three-step exponential decay. Bayesian model comparison showed the best performing models (M3 and M4) assumed 2 and 3 planning steps, while already assuming 1 planning step (M2) outperformed the no-planning model (M1 Δelpd > 880; **Fig. 2F,G**). The best-fitting models differed between subjects consistent with different planning depth : Subjects S and J were best fit by M4 (3-step planning), K by M3 (2-step planning), B and F by M2 (1-step planning), and R by M1 (no planning; **Fig. 2H**). The ordering of subjects by planning depth strategies matched the ordering of model-free estimated planning depth (**Fig. 2D**) with longer-to-shorter planning horizons ranking subjects from most to least planning into order S-J-K-B-F-R.

### Planning depth is predicted by knowledge of serial order

To understand which aspects of the sequence learning task predicted the difference in planning depth across subjects we extracted multiple performance metrics and used them as predictors of planning depth in linear mixed-effects models (LME’s). Consistent with planning reflecting serially choosing masked objects, forward planning on trials with masked objects was predicted by the metric ‘*maximum consecutive correct on masked trials*’ (MCC masked) (β = 0.95, p < .001) (**Fig. 3A**). Additionally, planning depth was predicted by the *proportion of trials subjects completed a sequence with zero errors* (perfect trial rate) (β = 1.99, p < .001), the *ratio of the minimum possible number of choices to the actual number of choices* across all trials (touch efficiency index) (β = 3.18, p < .001), as well as by the ‘*maximum consecutive correct overall*’ (MCC overall) (β = 0.94, p < .001) (**Fig. 3A**). These results suggest that better knowledge of the sequence is closely linked with planning. In contrast, learning speed metrics showed weak or non-significant effects, including the metrics *trial to first completion* (β = −0.07, p = .65), *trials needed to reach criterion performance* (β = 0.02, p = .78), and *cumulative learning errors* (β = −0.02, p = .39) (**Fig. 3A**). Session-level scatter plots confirmed that learning speed and cumulative errors had no within-subject relationship with planning depth once between-subject differences were accounted for (**Fig. S2**).

**Fig. 3.**
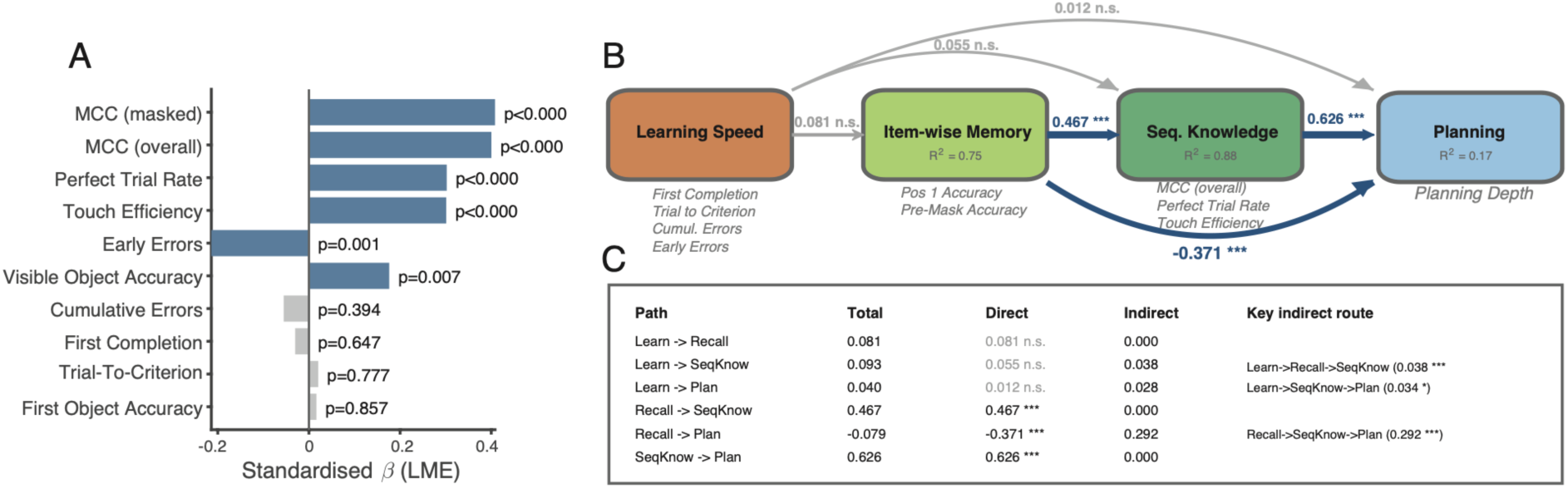
Structural path analysis linking memory quality to planning depth. (**A**) Linear Mixed Effects (LME) coefficients (β) for behavioral metrics predicting planning depth (all variables z-scored), with subject as random intercept. Blue bars indicate p < 0.05 (grey: n.s.). (**B**) Recursive structural path model with 4 constructs as nodes: Learning Speed → Item-wise Memory → Sequence Knowledge → Planning. In grey below boxes are the behavioral metrics defining it. Each node was regressed on all preceding nodes using LME’s with subject as random intercept. Grey numbers are coefficients for paths.Stars indicate sign. level. Sequence Knowledge predicted Planning (β = 0.63, p < .001). Item-wise Memory predicted Sequence Knowledge (β = 0.47, p < .001) but showed a negative direct path to Planning (β = −0.37, p < .001). Learning Speed had no direct effect on any downstream node. (**C**) Decomposition of total effect (*leftmost*) into direct (*middle*) and indirect (*rightmost*) paths effects for all links.

To test more directly for statistical dependencies among the behavioral metrics in predicting planning depth we used structural path analysis. For each subject, we grouped together the performance metrics that indexed how fast they learned the sequence (*Learning metrics*: e.g. trial to first completion), how well subject recalled objects (*Item Memory metrics*: e.g. pre-mask accuracy), and how well they recalled ordinal object transitions (*Sequence knowledge metrics*: e.g. overall proportion of consecutive correct choices). We entered these indices together with the estimated planning depth into a structural path analysis with the hypothesized processing sequence: Learning Speed → Item-wise Memory → Sequence Knowledge → Planning (**Fig. 3B**). The performance metrics at each endogenous node was regressed on all preceding nodes using linear mixed-effects models with subject as a random intercept. The analysis showed that planning depth was best predicted by Sequence Knowledge (β = 0.63, p < .001), while Sequence Knowledge was best predicted by Item-wise Memory (β = 0.47, p < .001) and Learning Speed had no direct effect on any downstream node (all p > .08). Path analysis allows decomposing the total effect into direct and indirect effects, which suggested that the direct link Item-wise Memory → Planning was near zero (β_total = −0.079) while Item-wise Memory contributed to a positive indirect path through Sequence Knowledge, i.e. Item-wise Memory → Sequence Knowledge → Planning (β = 0.292, bootstrap 95% CI [0.27, 0.45], p < .001) (**Fig. 3B,C**). This finding suggests that subjects who recalled individual items well and showed strong serial-order knowledge planned deeper, while subjects who recalled individual items well without building strong serial knowledge planned ahead less. This pattern of results indicates that item-level object memory is not sufficient and must be converted into ordered representations to support forward planning. Similar to Item-wise Memory, the Learning node that contained metrics on how fast subjects acquired a sequence, did not have a direct route to Planning (β_total = 0.04, n.s.), but a moderate indirect route via Sequence Knowledge (β_total = 0.034, p<0.05) (**Fig. 3C**).

### Planning forms a distinct cognitive factor

The analysis so far leaves open whether knowledge of a sequence is distinguishable from planning. To test how variations in planning depth relate to variations in other sequence performance metrics we used exploratory factor analysis (EFA) on session-level behavioral metrics. The Kaiser criterion supported a three-factor EFA solution explaining 83.3% of total variance (**Fig. 4A,B**). Factor 1 (54.0% variance) loaded on the metrics *maximum consecutive correct*, *perfect trial rate*, *touch efficiency* index, *position 1 accuracy*, and *pre-mask accuracy*. These metrics are consistent with labeling factor 1 ‘Sequence Memory Quality’. Factor 2 (18.3%) loaded on the metrics *first completion trial*, *trials to criterion*, *cumulative learning errors*, and *early error rate*, consistent with labeling factor 2 Learning Speed. Factor 3 (10.9%) loaded primarily on planning depth (loading = 0.75) and was labeled Planning. Planning thus formed a separate factor, separated from both memory quality and learning speed, indicating that it captures variance not explained by memory metrics alone. This conclusion is additionally supported by negative or near-zero correlations of the three latent factors (**Fig. 4C**).

**Fig. 4.**
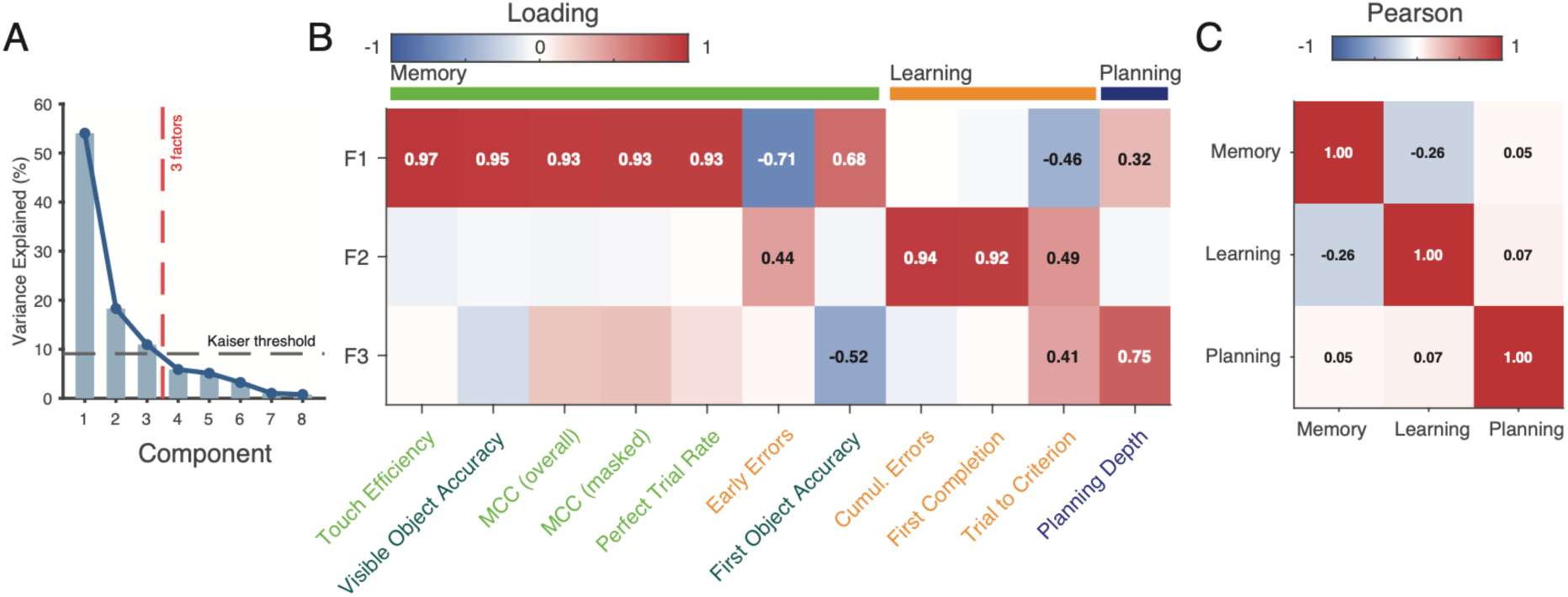
Sequence planning metrics separate memory, learning and planning factors. (**A**) Exploratory factor analysis (EFA) suggests 3 separate factors best describe the session-level behavioral metrics (within-subject z-scored, lower-is-better metrics inverted). Three factors explained 83.3% of total variance: Factor 1 (Sequence Memory Quality, 54.0%), Factor 2 (Learning Speed, 18.3%), and Factor 3 (Planning, 10.9%). (**B**) Factor loading heatmap after varimax rotation. Factor 1 loaded on sequence representation quality metrics (light green colored metric names). Factor 2 loaded on encoding speed metrics (orange). Planning depth loaded on its own third factor (loading = 0.75), distinct from both memory and learning factors. Dark green metric names are non-significantly contributing metrics. (**C**) Correlation matrix for the latent factors visualizes their distinctiveness.

### Planning and sequence memory is distinct from other canonical cognitive constructs

Our next question was whether planning and sequence learning are related to other higher-order cognitive processes. We tested the relationship of planning depth with four domain-general cognitive operations: inhibitory control, working memory, attention, and effort control (**Fig. 5**). Inhibitory control was measured with an antisaccade task that required inhibiting a pro-saccade response (**Fig. 5A**). All subjects showed lower accuracy and slower response times in the antisaccade than the prosaccade condition (**Fig. 5B,C**). Working memory capacity was measured with a continuous working memory updating task that measured how many objects subjects can add to their working memory before committing an error (**Fig. 5D**). Subjects updated on average 7.4 stimuli (SE = 0.15) before committing an error (using a 75% accuracy threshold) (**Fig. 5E**). Individual subjects had working memory capacities ranging from 6.04 (± 0.21 stimuli, monkey J) to 10.75 (± 0.31 stimuli, monkey F). When subjects made errors, they most likely erroneously chose an object that was added to working memory in earlier trials (**Fig. 5F**), consistent with a general memory decay. Selective attention was measured with a visual search task. Accuracy finding a target declined with increasing numbers of distractors when the distractor shared features with the target (high target-distractor similarity, slope = −0.021 ± 0.001, p < .001), while accuracy was rather independent of distractors when they differed from the target (slope: −0.002 ± 0.0005) (**Fig. 5H,I**). Effort control was measured with a task requiring subjects to choose between two options that differed in reward magnitude (1-5 drops of juice visualized using increasing number of tokens) and effort requirement (number of touches needed to inflate a ballon visualized as contours inside the balloons) (**Fig. 5J**). When obtaining a larger reward required additional effort, animals became progressively less likely to choose the larger-reward option as the effort difference increased (**Fig. 5K**; binomial GLM slope = -0.293 ± 0.006, p < .001). Conversely, animals became progressively more likely to choose the higher-effort option as the reward advantage of that option increased (**Fig. 5L**; binomial GLM slope = 0.320 ± 0.007, p < .001) (Fig. 5K,L). For the effort control metrics, choices were fit separately for each session using logistic regression with choice side as the outcome and reward difference and effort difference as predictors. The fitted reward and effort coefficients were used as session-level reward sensitivity and effort-cost weights, respectively.

**Fig. 5.**
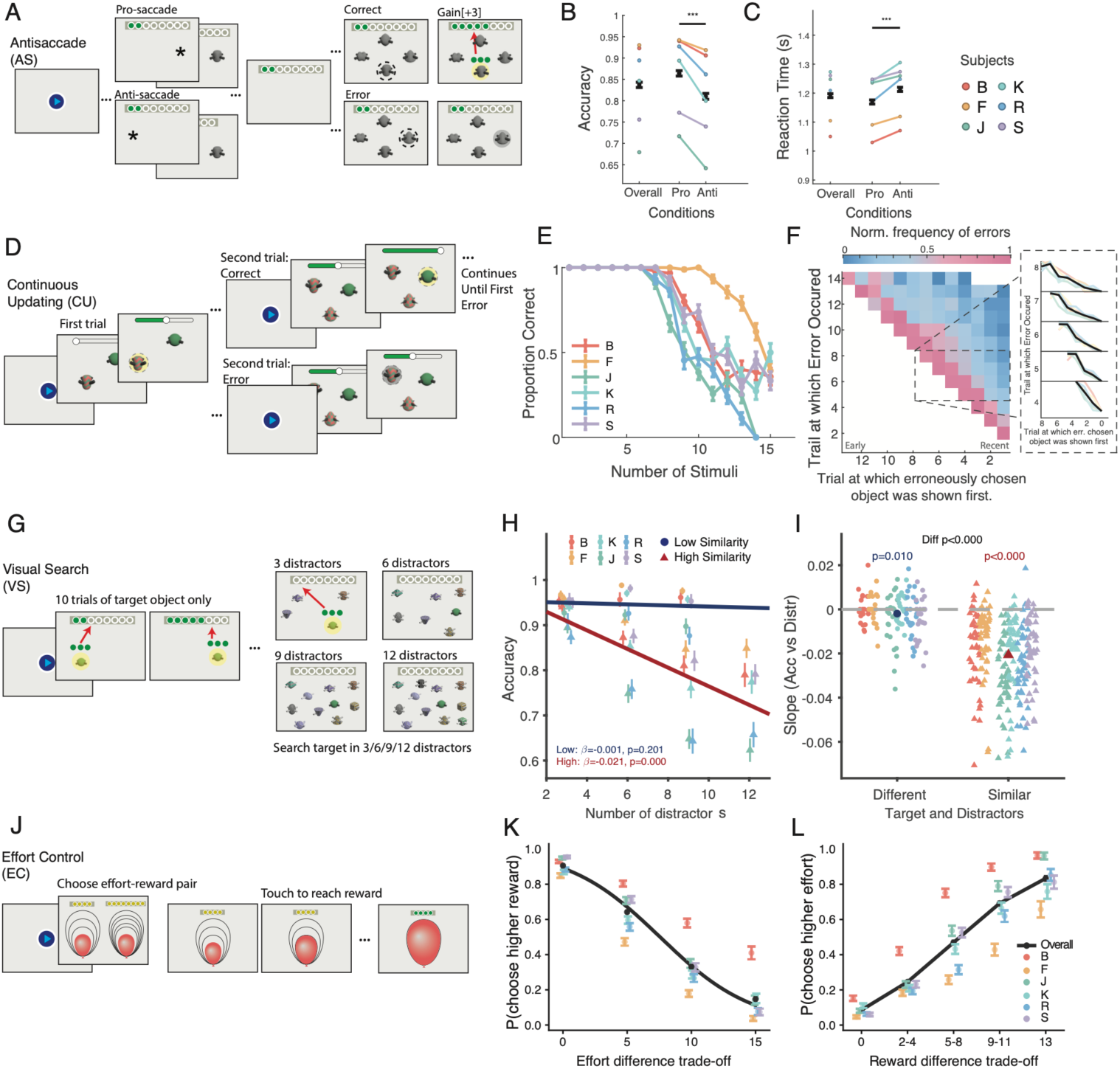
Robust assessment of inhibitory control, working memory capacity, attention and effort control. (**A**) The Antisaccade (AS) task flashed a peripheral cue and thereafter a target object at the cued location (pro-saccade) or the opposite location (anti-saccade). After a 0.5s delay, four objects appeared as response options and subjects selected the target to receive +3 tokens. (**B**) Accuracy was poorer for anti-saccade trials (0.81 ± 0.007 vs. 0.86 ± 0.006; paired t-test: t(273) = -12.67, p < .001). (**C**) Reaction times were slower for anti-saccade trials (1.21 ± 0.007 s vs. 1.17 ± 0.007 s; t(273) = -9.33, p < .001). (**D**) The Working Memory Updating (WM-U) task required subjects choosing an object they had not previously chosen. After every choice a new object was added. A block of trial ended when an error was made. (**E**) Across trials accuracy decreased with >5 objects for most subjects (diff. colors). (**F**) At each trial’s choice (*y-axis*), the most likely erroneously chosen object was from the immediate previous trial (x-axis). Inset: errors were most frequent for recently introduced objects. (**G**) The Visual Search (VS) task primed subjects with a target object and thereafter measured target detection with 3, 6, 9, or 12 distractors that were either similar or different to the target. (**H**) Highly similar distractors reduced accuracy, indexing a set-size effect of attentional interference. (**I**) Slopes of linear fits quantify robust set-size effects across sessions. (Slope in high-similarity condition: −0.021 ± 0.001, p < .001; low-similarity slope was near zero: −0.002 ± 0.0005, p = .010; high vs. low contrast: p < .001). (**J**) The Effort Control (EC) task required subjects to choose one of two effort–reward pairs. Reward magnitude was displayed as tokens, and effort level was indicated as number of balloon contours. When one option was chosen subjects inflated the chosen balloon by tapping the screen to cash out once inflated. (**K**) Subjects chose more likely the higher rewarded option (*y-axis*) when the demanded effort (*x-axis*: effort diff.) was lower. (**L**) Subjects chose the higher effort option when it earned higher reward (*x-axis*: reward diff.).

Planning and sequence learning performance might share variance with performance of these cognitive and motivational tasks. We used exploratory factor analysis (EFA) to test how shared performance variance was organized across metrics from all tasks. The results showed a robust six-factor solution (59.6% cumulative variance; Kaiser-Meyer-Olkin measure = 0.60, Bartlett p < .001; **Supplementary Results**) (**Fig. 6A**). Three factors separated metrics of the sequence learning task into a sequence performance and planning (factor 1), sequence item memory (factor 5) and learning of the sequence (factor 6). The three remaining factors combined metrics of the other tasks, grouping together visual search metrics (factor 2), working memory metrics together with antisaccade accuracy and effort control (factor 3), and effort control and working memory updating metrics (factor 4). Correlations of cross-factor pairs confirm that planning ability was distinct from other cognitive domains (**Fig. 6B,C**). The working memory factor 3 correlated with factor 5 (indexing sequence item memory) (Φ = 0.45, p = .004) but not with factor 1 (indexing planning and sequence memory), indicating that working memory abilities were linked to object level memory in the sequence task and only indirectly with planning (**Fig. 6C**). The selective attention factor 2 correlated with working memory factor 3 (Φ = 0.37, p = .008) and with the effort control factor 4 (Φ = 0.36, p < .001), but not with factor 1 that indexed planning. The dendrogram from hierarchical clustering of factor distances visually confirmed that the planning related factor 1 only indirectly related to other cognitive and motivational abilities (**Fig. 6C**).

**Fig. 6.**
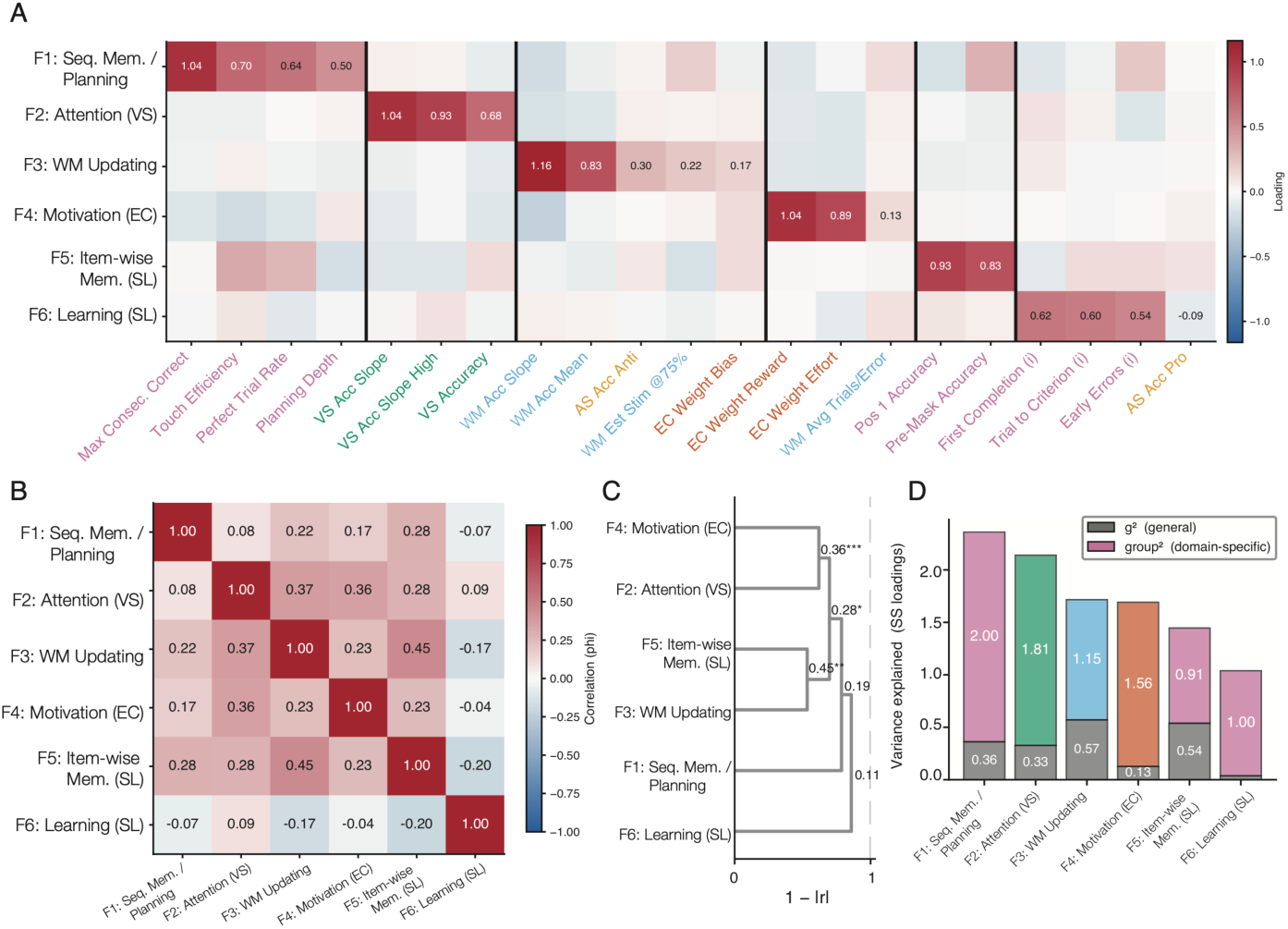
Planning and sequence learning form a cognitive factor separate from other cognitive domains. (**A**) Exploratory factor analysis across metrics of all five tasks suggest six factors (rows) that separate performance metrics (*x-axis*) signifying Sequence Knowledge (Factor 1), VS-based Attention (F 2), Working Memory Updating (F3), Effort Control/ Motivation (F 4), Seq. learning (SL) based Item Memory (F 5), and SL Learning Speed (F 6). Planning depth loaded with the Sequence Knowledge factor. Metrics were within-subject z-scored with lower-is-better metrics inverted. (**B**) Factor correlation matrix (Promax Φ). WM Updating (F3) and SL Item Memory (F5) correlated highest (Φ = 0.45, p = .004); most other pairs showed weak correlations (max |Φ| = 0.37). (**C**) Dendrogram of latent factor distances (*x-axis*: 1 − |r|, avg. linkage). Factor 4 (Effort control/Motivation) clustered with Factor 2 (VS Attention), and Factor 5 (SL Item Memory) with Factor 3 (Working Memory). (**D**) Results of bifactor model that assumed one common factor. Variance in each metric was partitioned into general factor (g) and six orthogonal domain-specific factors, shown as stacked bars of variance explained.

The availability of multiple task metrics made it possible to test how planning or the performance of any of the other measured cognitive domains are influenced by a common shared cognitive resource. We tested this with a bi-factorial model that assumes a common cognitive factor that subsumed performance variance shared across tasks in addition to task-specific variance. The overall explained common variance attributable to the general factor was 0.17, indicating that 17% of shared variance can be accounted for with a domain-general component. This finding shows that the overwhelming majority (83%) of variance shared across metrics is linked to task-specific factors. The amount of per-variable explained common variance for individual factors varied from 0.13 for factor 4 (indexing effort control) to 0.57 for factor 3 (indexing working memory) with a 0.36 common variance accounting for the planning factor 1 (**Fig. 6D**). These values are considered low and rule out that a common shared factor plays a major role accounting for planning performance of the sequence learning task, or for any of the other task (**Fig. S7**, **S8**).

## Discussion

We found that NHP’s plan ahead serial choices during object sequence performance (**Fig. 2B-D**). Planning ahead was a choice strategy that five of six subjects consistently engaged with as suggested by Bayesian modeling (**Fig. 2E-H**). Path analysis suggested that planning depth was predicted by the knowledge of the serial order, but only indirectly benefited from better object memory and faster learning of the serial order (**Fig. 3B,C**). Analysis of the shared and independent variance across performance metrics confirmed this conclusion by showing that planning depth – indexed as the likelihood of consecutive correct choices on masked objects - constitutes a distinct cognitive factor that was independent of sequence knowledge, object memory and learning abilities (**Fig. 4**). These findings quantify that planning ahead reflects a distinct cognitive operation that relies on an internal model of the serial order of objects. In support of forward planning as a distinct cognitive construct we found no or weak correlations with subject’s working memory capacity, attention, inhibitory control, and effort control.

### Planning ahead about two steps reflects a spontaneous choice strategy

We used serial object-order learning as the paradigm to probe planning ahead by introducing masks at variable times during sequential performance. At what stage the masking was introduced during the sequence was pseudorandomized in order to probe how subjects prospectively prepared their choices as opposed to using masks as a fixed cue that would have triggered a recall operation (Raby and Clayton, 2009). Five of the six subjects engaged in prospectively planning for at least two positions (**Fig. 2D**) with a model estimating planning horizons of 2-3 ordinal positions ahead (**Fig. 2E-H**) and factor analysis suggesting that successful planning ahead to choose masked object locations differs from the ability to choose unmasked objects (**Fig. 4**). This inferred planning depth likely does not reflect the limit of future planning, because the task paradigm did not enforce maximal planning requirements, but rather probed for planning either early or late during serial performance (with masks after the 1^st^, 2^nd^, or 3^rd^ choice) and included trials without planning requirements (no mask trials). With this mixture of trials subjects could have chosen low planning depth and still have received above-chance completion rate across trials, suggesting that planning ahead was a strategic choice subjects engaged with spontaneously. This feature of our task paradigm enabled analysis of the choice strategies (**Fig. 2H**) which showed that two subjects (S and J) showed a planning depth of three objects as their most preferred strategy, one subject (K) planned ahead 2 steps in the majority of sessions, two subjects (B and F) only occasionally showed planning depth above 1 step and one subject did not plan ahead (R). This planning variability is reminiscent on prior studies that trained nonhuman primates the serial order of images and then masked all objects after the first correct choice (Scarf et al., 2011). In this paradigm planning depth was two items after mask onset in most subjects while one subject reached above chance performance on the fourth item (Scarf et al., 2011). Similarly, three of four rhesus monkeys learning 5-object sequences have been found to anticipate the 4^th^ object in a sequence and rearrange it to the second position, providing evidence for position-specific coding of objects and some planning foresight of at least four objects (Wen et al., 2025b). The planning horizons of 2-4 in rhesus monkeys’ contrast with reports in chimpanzees, with some individuals being able to recall eight and more locations of learned digit symbols in the right numerical order after they were only briefly presented before being masked (Inoue and Matsuzawa, 2007). While there are likely genuine limits of considering future choice options in rhesus monkeys relative to chimpanzees (Raby and Clayton, 2009; Redshaw and Bulley, 2018), it is noteworthy that several chimpanzees showed prospective serial recall of learned numbers of only one or two items (Inoue and Matsuzawa, 2007; Muramatsu and Matsuzawa, 2023). These observations are consistent with the suggestion that planning abilities are influences by cognitive and motivational processes that vary across subjects, which we aimed to address in this study.

### Planning depth emerges from an internal knowledge model

We found with path analysis that knowledge of the object sequence was the strongest direct predictor of planning, while the ability to recall individual objects and the ability to efficiently learn object sequences only indirectly contributed to account for planning depth (**Fig. 3**). The close relation of knowing the serial order and perform the serial order when items were masked was corroborated in the larger-scale exploratory factor analysis that grouped together planning and sequence knowledge while separating them from item-wise memory, working memory, learning, attention, and effort control (**Fig. 6A,B**). These findings corroborate previous observations that the flexible mental re-arrangement of object order – which involves a form of planning ahead – was more likely when monkeys encoded serial object order using an abstract position code (Wen et al., 2025b). When monkeys learn serial relations among objects they initially use associative memory of item pairs, similar to chaining neighboring items (Terrace, 2005), but can develop the ability to represent the ordinal positions of objects consistent with a more abstract position code (Wen et al., 2025b), or by forming an abstract relational rule code (Sammeroff and Hampton, 2025). The more abstract representation of ordered information will facilitate planning operations. This suggestion resonates with computational frameworks that assume mental forward planning originates from internal models of the object order (Mattar and Daw, 2026). The internal models might be encoded as object-specific state transitions as in successor representation models, outcome probabilities of object positions as in inference models of planning (Botvinick and Toussaint, 2012; Solway and Botvinick, 2012), or as synaptic weights in multi-layer neural networks implementing a non-spatial cognitive map of the serial order of the object sequence (Behrens et al., 2018; Gershman, 2018; Whittington et al., 2022). All these computational architecture share the assumption that the ordinal object position within a sequence structure is represented in some abstract form to allow mental simulating choices of objects within that structure representation without executing the choices. Our finding of the importance of learned knowledge of the sequential order of objects is consistent with core assumptions of these models and provides predictions about the neuronal correlates of planning operations. In particular, our results suggest that neurons implementing planning ahead will recruit activity from neuronal populations representing the sequential structure with an abstract code instead of being tied to reward outcome related representations of stimulus-stimulus associations. Approaching these questions will involve identifying which computational planning framework best describes the behavioral pattern of planning we observed across monkeys.

### Planning ahead only indirectly relates to WM capacity, attention and effort control

We found that planning depth was not related to working memory capacity of individual subjects and was neither correlated with attentional abilities of interference control during visual search, with the efficiency to inhibit prepotent response in the antisaccade task or the willingness to exert effort to obtain higher reward in an effort control task (**Fig. 6**). These findings were unexpected and suggest that the ability to consider 2-3 future choices prior to the onset of masks – which is the basic metric reflecting planning in our study - is a distinct cognitive operation that is not reducible to a simple combination of working memory maintenance of choice options, covert attentional information sampling, suppressing non-favorable choice options and the availability of cognitive resources to engage in mental simulations without immediately acting. As a note of caution, the absence of correlations of these simpler cognitive operations and planning is a null finding and does not rule out that more complex planning tasks will recruit these other functions, or that dramatically larger sample sizes beyond the six NHP’s available for this study may reveal relationships among cognitive abilities. Beyond these limitations, our findings provide evidence for the absence of a direct relationship of working memory capacity and planning. Rather, working memory capacity was associated with better object level recall in the sequence learning task (latent factor correlation of Φ = 0.45, p = .004, **Fig. 6B**). According to the path analysis results object-level memory influences how well serial relationships can be learned (path link: 0.467, p<0.001) (**Fig. 3B**), suggesting that subjects’ working memory capacity positively influence object-level memory and thereby indirectly facilitates forming serial associations among objects. Our findings thus suggest that once working memory supports object level memory and enables sequence knowledge, subject will more likely engage in planning.

The outlined scenario is consistent with our main conclusion that planning is as a distinct operation, while at the same time outlining an indirect route for working memory capacity to facilitate planning operations. This hypothesized, indirect link of mental planning operations and working memory capacity reconciles various previous reports linking those processes. For example, in NHPs, mentally manipulating the serial order of objects correlate with lower (i.e. better) levels of short-term memory decay (Wen et al., 2025b), the self-ordering of items in working memory causally depend on the prefrontal cortex that also hosts object-location working memory (Petrides, 1995, 2005), the monitoring of an unfolding sequences is decodable in those prefrontal cortex areas that also support working memory (Yusif Rodriguez et al., 2024; Ahuja et al., 2026), and neuronal representations of serial ranks emerge as short-term memory buffer when subjects reverse items of that order (Tian et al., 2024). These examples suggest that working memory circuits are recruited when tasks involve an active manipulation of serial order. In our task paradigm, this mental manipulation primarily involved the spatial mapping of learned ordinal positions of objects to the spatial locations of objects on the screen prior to mask onset. There were no further mental manipulations needed than this prospective mapping of object order to spatial locations. A possible neural correlate of such a prospective mapping operation may be the neural firing in prefrontal cortex that predicts the execution of a sequence prior to the first choice (Averbeck and Lee, 2007). According to this scenario, planning ahead to correctly chose invisible masked objects in the right order may depend on the sequence memory for that order (see above) and the prospective recall of that memory via prefrontal cortex mechanisms that are distinct from working memory updating operations.

Taken together, our study documented that all but one NHP in our sample engaged in forward planning of 2-3 serial positions, and that planning was an independent cognitive construct that depended on the availability of good sequence knowledge, but only indirectly related to other cognitive operations. This pattern of results characterizes the cognitive architecture that underlies sequence learning and planning processes in NHP. Compared to humans NHP’s appear rather limited in their ability to plan and consider future states to guide behavior (Redshaw and Bulley, 2018). We believe that our study documents that these limitations will be gradual as inferred from NHP’s engaging spontaneously in future thinking about invisible objects using cognitive operations that are not easily reducible to more simple canonical cognitive functions.

## Materials and Methods

### Ethics statement

All animal and experimental procedures complied with the National Institutes of Health Guide for the Care and Use of Laboratory Animals and the Society for Neuroscience Guidelines and Policies and were approved by the Vanderbilt University Institutional Animal Care and Use Committee (IACUC) under approval number M1700198.

### Subjects and experimental procedures

Six male rhesus macaques (Subject B: 11 yrs/11.0 kg; Subject F: 13 yrs/11.9 kg; Subject J: 14 yrs/11.0 kg; Subject K: 13 yrs/11.3 kg; Subject R: 15 yrs/11.5 kg; Subject S: 11 yrs/11.5 kg) were used in this study. They performed the experimental tasks in their housing cages using cage-mounted touchscreen Kiosk stations (Womelsdorf et al., 2021), with example performance of three subjects shown in **Movie S1**. Visual display, behavioral response registration, and reward delivery were controlled by the Multi-Task Suite for Experiments (M-USE), an open-source video-engine based Unity3D platform integrated with a touchscreen, a video camera system, and reward delivery hardware (Watson et al., 2023).

Objects were multi-dimensional 3D rendered Quaddle 2.0 stimuli that varied in 10 feature dimensions (e.g., shape, color, body pattern, different arm orientations, the presence of a head, etc.), each with more than 10 possible feature values (Wen et al., 2025a). Novel sets of objects were generated for every new sequence by randomly assigning each object different features. The objects were generated using the software Blender and custom Python scripts that are freely available online (Wen et al., 2024). Object colors were chosen to be equidistant within the perceptually defined CIELAB color space. Objects were presented on an Elo 2094L 19.5″ LCD touchscreen with a refresh rate of 60 Hz and a resolution of 1,920 × 1,080 pixels, rendered at 2.9–4.2 × 2.4–4.7 cm on the screen. For each sequence, a new contextual background image was displayed. Context images were generated with DALL·E 2 (OpenAI) using text prompts for obtaining images from the category rocks. Color filters were applied to images to obtain a larger number of distinct context backgrounds, ensuring each color filter maintained a fixed luminance value of 50. The colors were selected by evenly spacing them around the CIELAB color wheel, ensuring a diverse range of hues. The brightness of the background images was adjusted to be ≤50% of the HSL scale.

### Behavioral paradigms

Subjects performed five tasks: Sequence learning and planning (SL), antisaccade (AS), effort control (EC), visual search (VS) and working memory updating (WM-U). SL, AS, EC, and VS were performed in the same order within the same experimental sessions every second weekday. The WM-U task was performed on the other days together with other tasks (set shifting, and visuospatial maze tasks) that are not considered in this report.

### Sequence learning and planning task

On each trial, five objects were presented at random locations at equal eccentricity relative to the center of the screen (**Fig. 1A**). Four of the five objects were assigned a unique ordinal position in a sequence. The fifth object was an irrelevant distractor object. Subjects learned to choose these four objects in the correct order by touching them with their fingers and receiving either positive feedback (a yellow halo and high-pitch sound) for correct choices or negative feedback (a transient grey halo and low-pitch sound) for incorrect choices. After an erroneous choice, subjects had to re-touch the last correct object in the sequence before searching for the next object. When a trial was completed or the maximal number of allowed errors were reached, objects were removed from the screen and a new trial began with the objects displayed at new random locations equidistant from the center of the display.

For each correctly chosen object, the slider position of a progress bar on top of the screen stepped forward. Successful completion of a sequence completed the progress bar and resulted in a fluid reward provided though a sipper tube in front of the monitor. When a sequence was not completed, no fluid reward was given and the progress bar was reset for the subsequent trial. The progress bar provided additional visual feedback about whether the subject’s choice was correct or erroneous and how many choices remained before sequence completion and reward.

Each session contained five sequences of 20 trials each (**Fig. 1C**). Within each sequence, trials 1 through 5 served as unmasked learning trials in which all objects were visible on every trial. Trials 6 through 20 introduced a masking manipulation: on a given trial, objects at and beyond a randomly assigned ordinal position were hidden from view, requiring the subject to rely on memory to select the correct object. The masking onset position varied across trials, with masking beginning at ordinal position 2, 3, or 4, each assigned with approximately 30% probability, while the remaining approximately 10% of trials in the masked phase remained fully visible as unmasked control trials (**Fig. 1B**). This design allowed us to assess both how well subjects learned each four-item sequence and how accurately they could recall masked items at different ordinal positions.

### Antisaccade task

The antisaccade (AS) task measured inhibitory control (**Fig. 5A**). Subjects started a trial by touching a trial-start button, after which a cue appeared at a peripheral location. A target object was then briefly presented either at the cue’s location (pro-saccade condition) or at the opposite location (anti-saccade condition). After a brief delay, four objects appeared as response options and subjects had to select the target object. Correct selections were rewarded with three tokens. A total of 274 sessions were included in the AS analysis.

### Working memory updating task

The working memory updating (WM-U) task measured WM capacity and updating abilities (**Fig. 5D**). The first trial presented two objects; each subsequent trial added one additional object to the display. Subjects had to choose an object they had not chosen in any previous trial to receive fluid reward and progress a slider position. The task continued until the subject made an error, after which the current trial ended and a new trial began. This task required subjects to maintain and update a growing set of items in working memory, with accuracy reflecting the number of items that could be tracked reliably. A total of 271 sessions were included in the WM-U analysis.

### Visual search task

The visual search (VS) task measured selective attention (**Fig. 5G**). Subjects first performed 10 trials in which they repeatedly chose a single target object to familiarize them with its identity. They then searched for this target among 3, 6, 9, or 12 distractors. The target-distractor similarity was manipulated across blocks by varying the number of object features shared between targets and distractors: in the low-similarity condition, distractors had unique features and were perceptually distinct from the target, while in the high-similarity condition, distractors shared >1 feature with the target. Accuracy was measured as a function of the number of distractors and the similarity condition. A total of 271 sessions were included in the VS analysis.

### Effort control task

The effort control (EC) task indexed motivation to exert effort for reward (**Fig. 5J**). On each trial, subjects chose between two options that differed in reward magnitude and effort requirement. Reward magnitude was indicated by the number of reward tokens displayed on top of each option and which corresponded to different number of fluid drops provided as primary reward after trial completion. Effort was indicated by different numbers of contour lines within the balloons, from small to large, representing the number of screen taps required to fully inflate the balloon and collect the reward. Subjects made their choice by selecting one option and then completing the required tapping to receive the reward. When subjects chose the option with less effort (less contours) there was a delay introduced between trials that was as long as it would have taken to inflate the alternative balloon with more contours. The adjustment of this inter-trial delay ensured that both effort options had similar wait times till the next trial started, so that effort corresponded to manually touching the ballon and not to temporal costs until the next trial started. A total of 274 sessions were included in the EC analysis.

### Behavioral metrics

All metrics of the sequence and learning task were computed per sequence and then averaged across the five sequences within a session to yield one value per session per subject. Subject means and standard errors were obtained across sessions. Unless noted otherwise, linear mixed-effects models (LME) with subject as a random intercept were used for statistical inference. Metrics were organized into four categories: learning speed, item-wise memory, sequence knowledge, and planning.

*Learning speed* metrics included: the first completion trial (FCT), defined as the trial number on which the subject first completed the full four-item sequence correctly; the trials to criterion (TTC), defined as the first trial on which the subject completed the sequence on two consecutive trials, which avoided considering a single completion as reflecting stable learning; cumulative learning errors (CLE), which counted the total number of incorrect choices accumulated from trial 1 up to and including the trial of first completion (re-touches of the last correct objects after an error were not counted as they reflected task rule; if the sequence was never completed within 20 trials, this metric was considered missing to avoid upward bias); and the early error rate, computed as the proportion of incorrect touches during the first five unmasked trials.

*Item-wise memory* metrics included: accuracy at the first ordinal position across trials 6 through 20 (position 1 accuracy), which reflects the recall of an object at the first position that was never masked; and pre-mask accuracy, quantified as mean accuracy at ordinal positions preceding the mask onset on masked trials, providing a second measure of item-level recall uncontaminated by the mask manipulation.

*Sequence knowledge* metrics included: the touch efficiency index (TEI), computed as the ratio of the minimum possible number of object choices to the actual number of task-relevant choices across all trials in the sequence (a value of 1.0 corresponds to perfect performance with no errors on any trial); the maximum consecutive correct (MCC) selections per trial, defined as the longest streak of consecutive correct choices within a trial regardless of masking condition; the perfect trial rate (PTR), defined as the proportion of trials after the first completed trial on which the subject completed a sequence with zero errors; and correct state transition probabilities, measuring the probability of advancing from each ordinal state to the next on the following touch, separately for each of the four transitions (state 0→1, state 1→2, state 2→3, and state 3→4), computed on no-mask trials (trials 11–20) and compared against forward-looking chance (1 divided by the number of remaining objects at each state).

### Planning depth estimation

To assess how far ahead subjects planned during masked trials, we computed separately for each masking condition the proportion of correct choices for each of the four ordinal positions of trials after subjects had learned the sequence, i.e. after they successfully completed two consecutive trials (**Fig. 2A**). At each ordinal position, we compared accuracy against a forward-looking chance level that accounted for previous correct selections within the trial (1/N remaining objects). To quantify choice accuracy relative to the position at which the mask was presented first, we aligned all masked trials by the position at which masking began and computed the difference between actual accuracy and forward-looking chance at each relative position (**Fig. 2B**). Grand-mean accuracy differences were tested at each relative position using LME models with subject as a random intercept, testing whether the intercept was significantly above zero. To estimate a continuous measure of planning depth for each session, we fitted a linear function to the accuracy-minus-chance profile at relative positions 1, 2, and 3 from mask onset and found the interpolated zero-crossing of this profile (**Fig. 2C,D**). If all three positions remained above chance, planning depth was set to 3; if the first position was at or below chance, depth was set to 0. This yielded a continuous planning depth estimate per session, clipped to the range [0, 3].

### Controlling for short-term memory decay of objects since mask onset

To test whether above-chance performance after mask onset could be explained by variation in retention delay, we calculated the elapsed search time since the final unmasked selection within each trial. For the first position after mask onset, each masked condition was restricted to the shared 10th-90th percentile delay window across conditions (0.662-3.084 s). Accuracy minus forward-looking chance was then recomputed within this matched window using LME models with subject as a random intercept (**Fig. S1I**).

### Error type classification

To characterize the nature of errors, we classified each errorneous choice into one of two categories. Rule-abiding errors were incorrect choices on one of the sequence-relevant objects (i.e. choosing a sequence object out of order). Rule-breaking errors were choices that violated the task structure, such as failing to re-choose the last correct object after an error. We also classified errors by whether the chosen object was a sequence-relevant object selected out of order (sequence error) or the sequence-irrelevant distractor (distractor error). The proportion of each error type was computed across trials 11–20 and compared using LME models with error type as a fixed effect and subject and session as random effects.

### Strategy comparison using Bayesian modeling

To formally compare planning strategies, we fitted four Bayesian models to the masked trials accuracy data (**Fig. 2E-H**). Data were aggregated per subject × relative-mask-position × ordinal-position cell so that the per cell chance level (1/N remaining objects) was exact rather than averaged. Let cell *i* belong to subject *j(i)*, with *n_i* trials and *k_i* correct choices. All four models shared a binomial likelihood, *k_i* ∼ Binomial(*n_i*, *p_i*), where *p_i* = clip(*c_i* + Δ*_i*, 0.01, 0.99) and *c_i* is the forward-looking chance for that cell. The models differed only in the accuracy boost Δ*_i*: Model M1 (no planning) set Δ*_i* to a small subject-level offset around zero; Model M2 (one-step) applied a per-subject boost *b_j* only at the first masked position; Model M3 (two-step) applied an exponentially decaying boost *b_j* · exp(−*r_i* / *τ_j*) across the first two masked positions; and Model M4 (three-step) applied the same exponential decay across all three masked positions. Subject-level parameters (*b_j*, *τ_j*) were drawn from population-level priors (partial pooling): *b_μ* ∼ Normal(0.15–0.20, 0.10), *b_σ* ∼ HalfNormal(0.10), *b_j* ∼ Normal(*b_μ*, *b_σ*); for S3 and S4, *τ_μ* ∼ HalfNormal(2.0), *τ_σ* ∼ HalfNormal(1.0).

Models were fitted using the No-U-Turn Sampler (NUTS) implemented in PyMC (v5), with 2 chains, 500 tuning steps, and 1,000 draws per chain (2,000 posterior samples total). Target acceptance rate was set to 0.80 for M1 and M2 and 0.90 for M3 and M4 to accommodate the hierarchical decay parameter. Convergence was confirmed by R-hat = 1.00 for all subject-level parameters and effective sample sizes (ESS_bulk) ≥ 600, with no divergent transitions. Model comparison was performed using the WAIC (Widely Applicable Information Criterion) (**Fig. 2F**) and Pareto-smoothed importance sampling Leave-One-Out cross-validation (PSIS-LOO; **Fig. 2G**). Both metrics were expressed as expected log predictive density (elpd), with higher values indicating better fit. Subject-level strategy assignments were determined by selecting the model with the highest elpd for each subject.

### Univariate predictor screening

To identify which behavioral metrics predicted planning depth, we conducted univariate predictor screening using separate LME models for each z-scored predictor against planning depth with subject as a random intercept (**Fig. 3A**).

### Structural path analysis

To test the causal architecture linking learning, memory, and planning, we conducted an observed-variable recursive path analysis with four constructs ordered by their hypothesized processing sequence: Learning Speed → Item-wise Memory → Sequence Knowledge → Planning. For each construct, a score was computed using the mean of the z-scored behavioral metrics that defined the composite score. Learning Speed comprised the metrics first completion trial, trial to criterion, cumulative learning errors, and early error rate (reverse-scored so that higher values indicate faster learning). Item-wise Memory comprised position 1 accuracy and pre-mask accuracy. Sequence Knowledge comprised maximum consecutive correct, perfect trial rate, and touch efficiency index. Planning comprised planning depth.

Each endogenous node was regressed on all preceding nodes simultaneously using LME models with subject as a random intercept, yielding direct path coefficients for all six forward links. Total effects were decomposed into direct and indirect components using path tracing rules (**Fig. 3B,C**). Confidence intervals for indirect effects were obtained via cluster bootstrap (2,000 resamples at the subject level), resampling entire subjects with replacement and re-estimating all path coefficients at each iteration. Bootstrap p-values were computed as twice the proportion of bootstrap samples in which the indirect effect crossed zero.

### Within-task exploratory factor analysis

To determine whether planning reflects a distinct cognitive process or emerges from the quality of underlying memory representations, we performed exploratory factor analysis (EFA) on session-level behavioral metrics from the sequence learning and planning task (**Fig. 4**). Eleven metrics were included after applying the following screening criteria (**Fig. S4**): split-half reliability greater than 0.70, non-redundancy (pairwise correlation below 0.80, retaining one of highly correlated pairs), and adequate coverage (not missing in most sessions). We computed the Kaiser-Meyer-Olkin (KMO) measure of sampling adequacy for each metric (**Fig. S4C**) and communalities from the fitted EFA (**Fig. S4D**). The number of factors was determined by the Kaiser criterion (eigenvalue > 1). Varimax rotation was applied to obtain interpretable loadings. The latent factor correlation matrix was computed to assess inter-factor relationships (**Fig. 4C**).

### Cross-task exploratory factor analysis

To examine whether the cognitive processes underlying planning and sequence learning are shared with other tasks, we conducted exploratory factor analysis (EFA) with oblique (Promax, power = 4) rotation across session-level metrics from all five tasks (**Fig. 6A**). Metrics were screened using the same reliability and non-redundancy criteria as the within-task EFA, yielding 21 variables across 154 paired-session observations. The number of factors was determined by parallel analysis combined with inspection of the scree plot and factor interpretability. The Kaiser-Meyer-Olkin measure of sampling adequacy and Bartlett’s test of sphericity were computed to confirm the suitability of the data for factor analysis. The six-factor solution was validated against alternative rotation methods (varimax; **Fig. S5A**) and alternative factor numbers (five and seven factors with Promax rotation; **Fig. S5B,C**).

At the individual metric level, we computed pairwise cross-task correlations between each sequence learning metric and each metric from the four concurrent tasks (**Fig. S3A**). We also computed RV coefficients (a multivariate generalization of R²) between each group of sequence learning metrics (Learning Speed, Item-wise Memory, Sequence Knowledge, Planning) and each concurrent task (**Fig. S3B**). To assess whether cross-task associations reflected stable between-subject trait differences or session-to-session variability, we repeated the correlation analysis after averaging session data within subjects (**Fig. S7**).

### Inter-factor correlation and hierarchical clustering

An inter-factor correlation matrix was computed to assess associations between latent factors (**Fig. 6B**). To identify the hierarchical structure among factors, we computed pairwise distances as 1 − |r| and applied hierarchical clustering with average linkage and displayed the result as a dendrogram (**Fig. 6C**). Factor-pair associations were tested using two-sided Pearson correlations between factor scores.

### Bifactor analysis

To test whether task performance reflected a shared general ability, we applied a bifactor model via Schmid-Leiman (SL) orthogonalization (**Fig. 6D**). This procedure transforms the oblique first-order Promax solution into one general factor (*g*) plus orthogonal residual group factors. A single higher-order factor was fitted to the first-order factor correlation matrix Φ by maximum-likelihood factor analysis, yielding higher-order loadings *B_j* (the loading of each first-order factor *j* on *g*) and uniquenesses *U²_j* = 1 − *B_j*². The original first-order pattern matrix *P* was then decomposed into a general component (*g* loadings = *P* · *B*) and residual group components (group loadings = *P* · diag(√*U²*)), such that each variable’s communality partitioned additively into general and group contributions: *h²_i* = *g_i*² + Σ*_j* *G_ij*². The overall explained common variance (ECV) was computed as the ratio of the sum of squared *g* loadings to the total sum of squared loadings (general plus group). Per-variable ECV was computed for each metric to assess how much of each metric’s communality was attributable to *g* versus domain-specific factors (**Fig. S6A,B**). An ECV ≥ 0.60 is conventionally taken as evidence for essential unidimensionality. Higher-order factor extraction was performed using the FactorAnalyzer library in Python with maximum-likelihood estimation, one factor, and no rotation applied to the first-order correlation matrix.

### Statistical analysis

All statistical analyses were conducted in MATLAB R2024b (MathWorks) and Python 3.9.12. The Python environment included NumPy 1.26.4, pandas 2.0.0, SciPy 1.12.0, and scikit-learn 1.4.2. Linear mixed-effects models were fitted with MATLAB’s fitlme function, and generalized mixed-effects models for binary outcomes were fitted with fitglme. Exploratory factor analyses were performed with MATLAB’s pca and rotatefactors functions and the Python FactorAnalyzer package, as specified for each analysis. Multiple comparisons were corrected using Bonferroni correction unless otherwise stated. Bootstrap confidence intervals used 2,000 resamples with cluster resampling at the subject level. All p-values were two-tailed.

## Acknowledgements

This work was supported by the National Institute of Mental Health of the National Institutes of Health under Award Number R01MH129641 (TW). The content is solely the responsibility of the authors and does not necessarily represent the official views of the National Institutes of Health.

## Data, Materials, and Software Availability

The data and analysis code supporting this study will be available in a public repository on Zenodo upon publication. All other data are included in the article and/or supporting information.

## Author Contributions

A.N. and X.W. performed research; X.W. analyzed data; S.D. contributed to the task software platform; X.W., P.T., T.W. wrote and edited the paper; T.W. supervised the project.

