## Supplementary Material for "Planning in Nonhuman Primates Emerges from Structure Knowledge and is Distinct from Attention, Working Memory, Effort Control, and Learning"

**Content:**

**Supplementary Methods**

**Supplementary Results**

**Supplementary Figures S1-S8**

**Supplementary Methods**

**Search time analysis.** Search time was defined as the time from trial onset (or from the previous touch) to each touch. We computed the distribution of search times at masked positions separately for the first, second, and third masked positions (Fig. S2A,B). To test whether faster responses were associated with higher accuracy, we split trials into the fastest 25% and slowest 25% by search time and computed the delta accuracy (accuracy minus forward-looking chance) at each masked position for each speed quartile (Fig. S2C).

**Supplemental Results**

**Analysis of choice reaction times and the effects of a speed-accuracy tradeoff**

When trials were split by response speed, faster responses were associated with higher accuracy that remained above chance at all masked positions, while slower responses also stayed above chance but with lower accuracy (**Fig. S2A-C**). This pattern suggests that faster retrieval reflected stronger pre-loaded representations, rather than a speed-accuracy trade-off.

**Exploratory factor analysis of planning and cognitive and motivational domains**

We used exploratory factor analysis with oblique (Promax) rotation across metrics from the four tasks together with metrics of the planning and sequence learning task. The results support a six-factor solution (59.6% cumulative variance; Kaiser-Meyer-Olkin measure = 0.60, Bartlett p < .001) (**Fig. 6A**): Factor 1 loaded on maximum consecutive correct, touch efficiency index, perfect trial rate, and planning depth from the sequence learning task and labeled SL; Factor 2 loaded on visual search accuracy metrics (label VS); Factor 3 loaded on working memory updating accuracy slopes (label WM); Factor 4 on effort control weights (label EC); Factor 5 on position 1 and pre-mask accuracy of the sequence learning task (label SL Item Memory); and Factor 6 loaded on first completion trial, trial to criterion, and early errors (LS Learning Speed). Inhibitory control metrics did not show latent loadings. Planning depth loaded with the Sequence Knowledge factor (loading = 0.50), consistent with the within-task finding that planning and sequential knowledge share a latent dimension. The six-factor solution was robust to alternative rotation methods and factor numbers (**Fig. S5A–C**). At the individual metric level, pairwise cross-task correlations were generally weak, with only a few significant pairs at moderate p-values (**Fig. S3A**). RV coefficients between each SL metric group and each concurrent task were similarly small (**Fig. S3B**). These cross-task associations appeared to be largely subject-specific: when session data were averaged within subjects, the correlations were no longer visible (**Fig. S7**), suggesting that the metric-level associations were driven by session-to-session variability rather than stable between-subject trait differences.

**Supplemental Figures S1-S8**

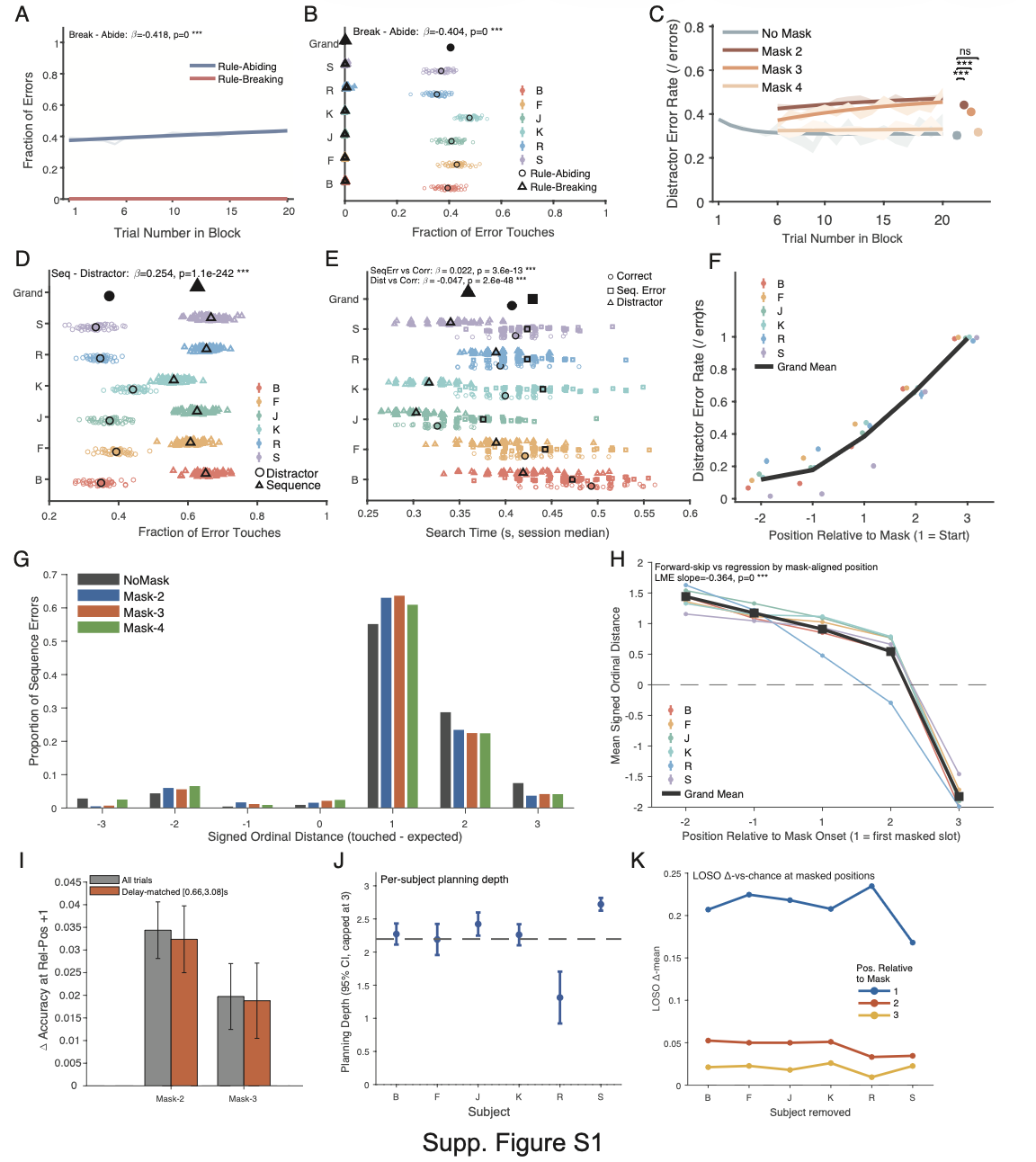

**Supplementary Figure S1.** **Sequence Errors Were Structured and Planning Measures Survived Control Analyses.**

(A) Proportion of rule-abiding and rule-breaking errors across trial position (1–20), fitted with exponential decay curves with SEM bands. Rule-abiding errors are incorrect touches on objects that belong to the sequence but were selected out of order; rule-breaking errors are touches on objects not in the sequence at all. Trials 11–20 mean: rule-abiding = 0.419 ± 0.004, rule-breaking = 0.002 ± 0.0003 (LME: β = −0.418, p < .001).

B) Per-session fraction of rule-abiding versus rule-breaking errors for each subject, shown as horizontal jittered dot plots. All subjects showed rule-abiding errors comprising 35–48% of errors (R: 0.35; K: 0.48) and rule-breaking errors near zero (all ≤ 0.007; LME: β = −0.404, p < .001). R had the highest rule-breaking rate (0.007), though still negligible. (C) Distractor error proportion (fraction of errors that were distractor touches, i.e., touching the non-sequence object) across trial position for each masking condition, fitted with exponential decay. Trials 11–20 (LME with Bonferroni correction): no-mask = 0.302 ± 0.010; mask 2 = 0.441 ± 0.005 (p < .001 vs. no-mask); mask 3 = 0.410 ± 0.005 (p < .001); mask 4 = 0.317 ± 0.008 (p = .29, n.s.). Distractor errors were elevated when early items were masked but not when only the last item was masked. (D) Per-session fraction of distractor versus sequence-related errors for each subject, shown as horizontal dot plots. Approximately one third of errors were distractor touches (0.373 ± 0.003) and two thirds were wrong-order sequence touches (0.627 ± 0.003; LME: β = 0.254, p < .001). Per-subject distractor fraction ranged from 0.334 (S) to 0.441 (K). (E) Search time (session median, in seconds) for correct touches, sequence errors, and distractor errors, shown as horizontal per-subject dot plots. Correct touches: 0.407 ± 0.003 s; sequence errors: 0.429 ± 0.003 s; distractor errors: 0.360 ± 0.003 s. Sequence errors were significantly slower than correct touches (β = +0.023 s, p < .001) and distractor errors were significantly faster (β = −0.048 s, p < .001), suggesting distractor errors reflect impulsive responses rather than deliberative mistakes. (F) Distractor error rate (fraction of errors that were distractor touches) aligned to ordinal position relative to mask onset (relative position −2 to +2), on post-criterion masked trials. Per-subject jittered session dots with grand mean and SEM band. Monotonic increase: 0.118 ± 0.007 (relative position −2), 0.178 ± 0.008 (−1), 0.385 ± 0.009 (0), 0.668 ± 0.007 (+1), 0.991 ± 0.003 (+2). Once items were hidden, errors became increasingly likely to be distractor touches. (G) Forward-skip error topology by mask condition. Mean signed ordinal distance (d = touched position − expected position) for sequence errors, split by masking condition. Positive values indicate forward skips; negative values indicate backward regressions. Errors were biased forward in every condition (all p < .001): no-mask = 1.172 ± 0.013, mask-2 = 1.057 ± 0.010, mask-3 = 1.066 ± 0.012, mask-4 = 0.965 ± 0.014. The forward bias was attenuated as more items were masked (LME slope = −0.045, p < .001). (H) Forward-skip error topology aligned to mask onset. Mean signed ordinal distance at positions relative to mask onset (relative position −2 to +2), on post-criterion masked trials. Forward bias was strongest before mask onset (relative position −2: 1.442 ± 0.010; −1: 1.173 ± 0.010) and decayed monotonically across masked positions (0: 0.909 ± 0.013; +1: 0.544 ± 0.017), becoming backward at relative position +2 (−1.826 ± 0.033). LME slope = −0.364, p < .001. (I) Delay-matched working-memory control. Δ-accuracy (accuracy − forward chance) at the first masked position, restricted to a common retention-delay window [0.662, 3.084 s] across mask conditions, compared with the unmatched estimate. The above-chance accuracy survived delay matching at mask-2 (Δ = 0.032, p < .001) and mask-3 (Δ = 0.019, p = .024). Mask-4 yielded no trials within the matched window and could not be evaluated. (J) Leave-one-subject-out planning-depth robustness. Each subject was removed in turn and group-mean planning depth and per-relative-position Δ-accuracy were recomputed on the remaining five subjects. All six leave-one-out group means remained between 2.09 and 2.37, and the above-chance accuracy at the first masked position was significant in every subsample (all p < .04; max p = .031 when S was removed). Per-subject planning depth (mean ± 95% CI) is also shown. (K) Trial-state-controlled conditional dependency. The conditional chaining analysis from Fig. 2E was refit with a trial-level random intercept to absorb trial-state variance. The prior-correct effect survived this control (β = 0.142, p < .001). A mask condition × prior-correct interaction was not significant (p = .42), and the chaining effect was present in both unmasked (β = 0.112, p < .001) and masked (β = 0.154, p < .001) trials, indicating that sequential dependency was a general property of performance rather than an artifact of trial-level fluctuations. N = 6 subjects, 274 sessions.

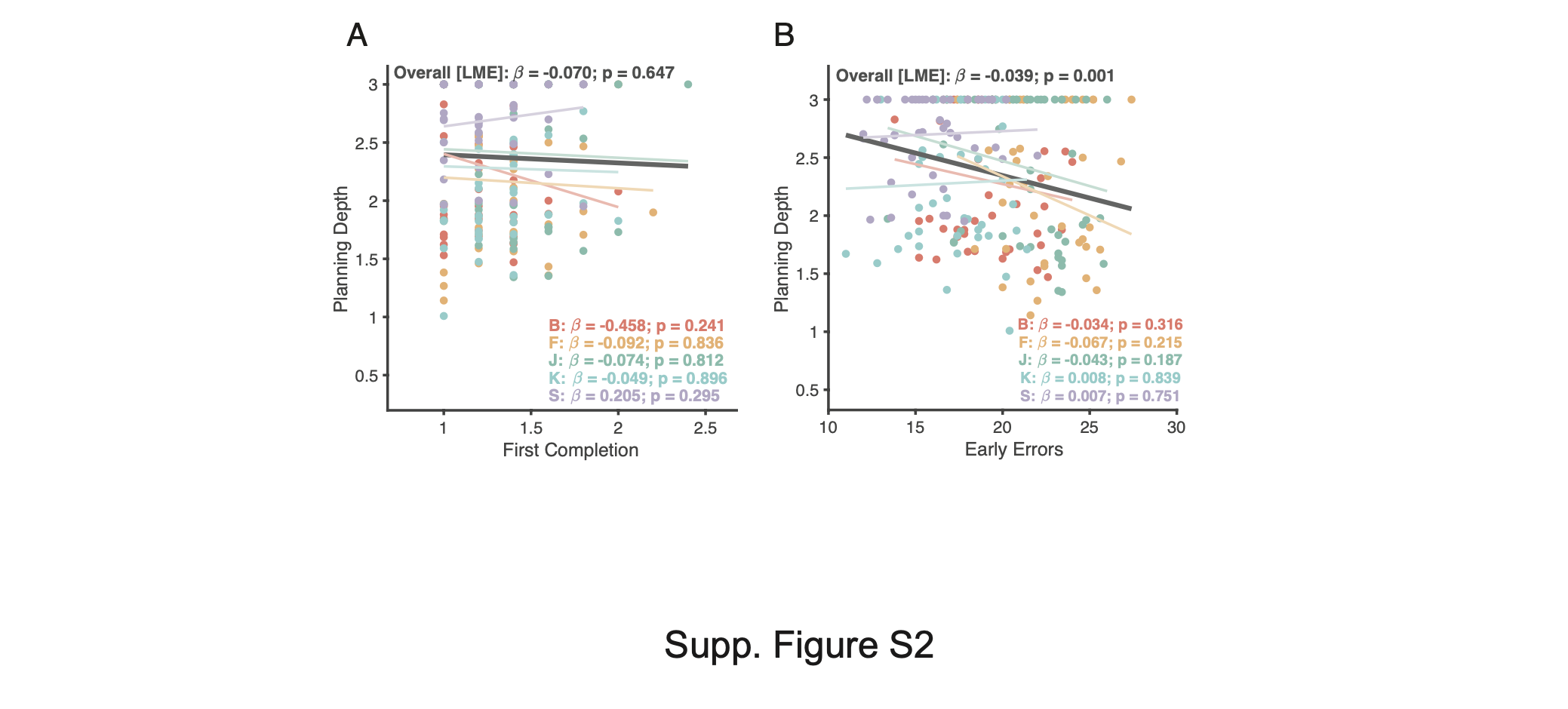
**Supplementary Figure S2. Planning Depth Was Unrelated to Early Learning Speed.**

(A) Session-level scatter plot of first completion trial versus planning depth, with subject identity indicated by color. LME regression line with β and p annotated. (B) Session-level scatter plot of cumulative learning errors versus planning depth, with subject identity indicated by color. LME regression line with β and p annotated.

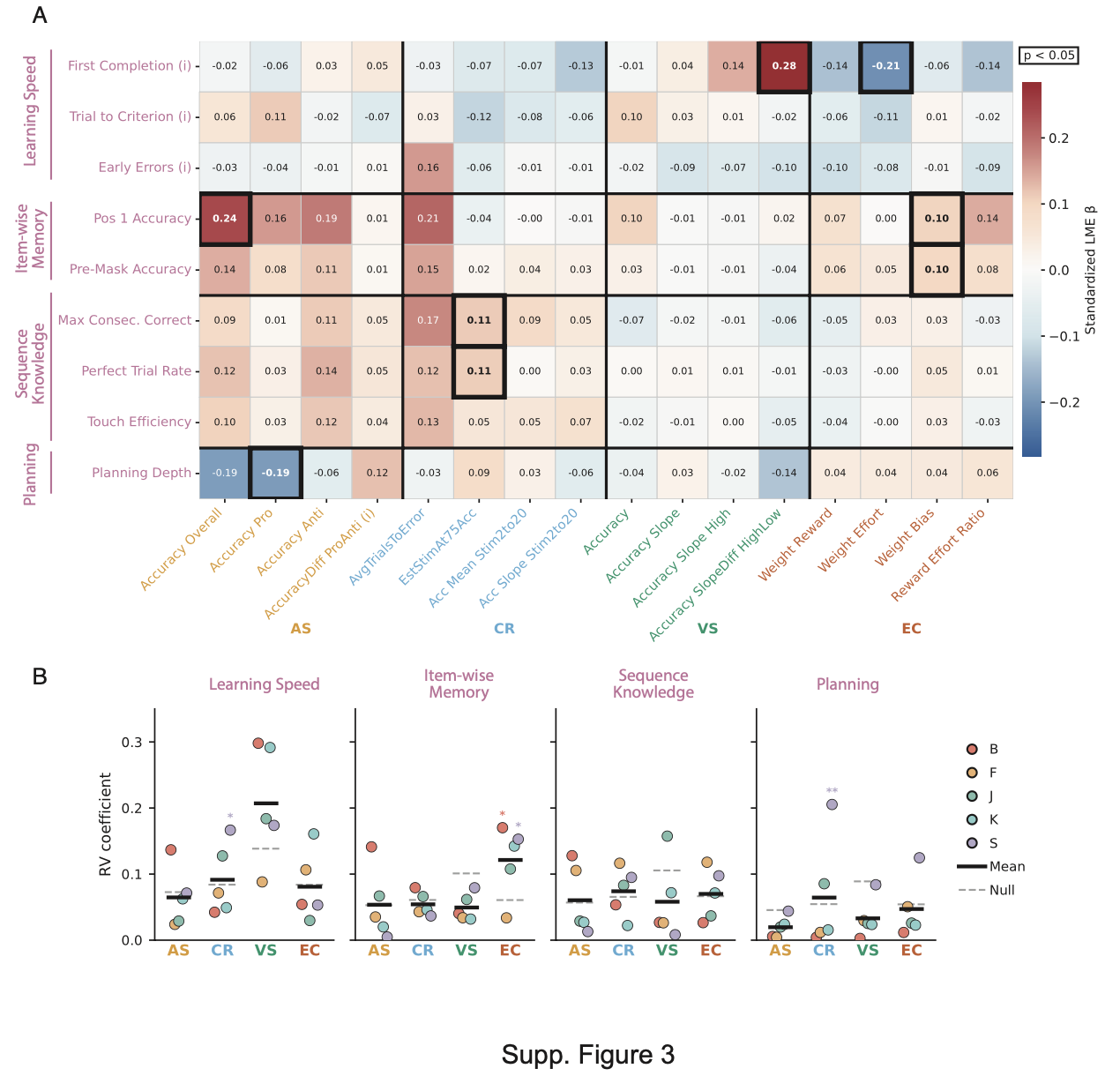
**Supplementary Figure S3. Sequence Learning Metrics Had Limited Cross-Task Associations.**

(A) Correlation matrix of all sequence learning metrics against metrics from the four concurrent tasks (antisaccade, continuous recognition, visual search, effort control). Pearson correlations computed on session-level within-subject z-scored metrics. Significant pairs (uncorrected p < .05) are indicated by outlined boxes. (B) RV coefficient (Escoufier) between each SL metric group (Learning Speed, Item-wise Memory, Sequence Knowledge, Planning) and each concurrent task metric matrix, computed per subject. The RV coefficient measures the overall association between two sets of variables, ranging from 0 (no association) to 1 (perfect association). Bars show subject-level RV values. N = 6 subjects, 274 sessions.

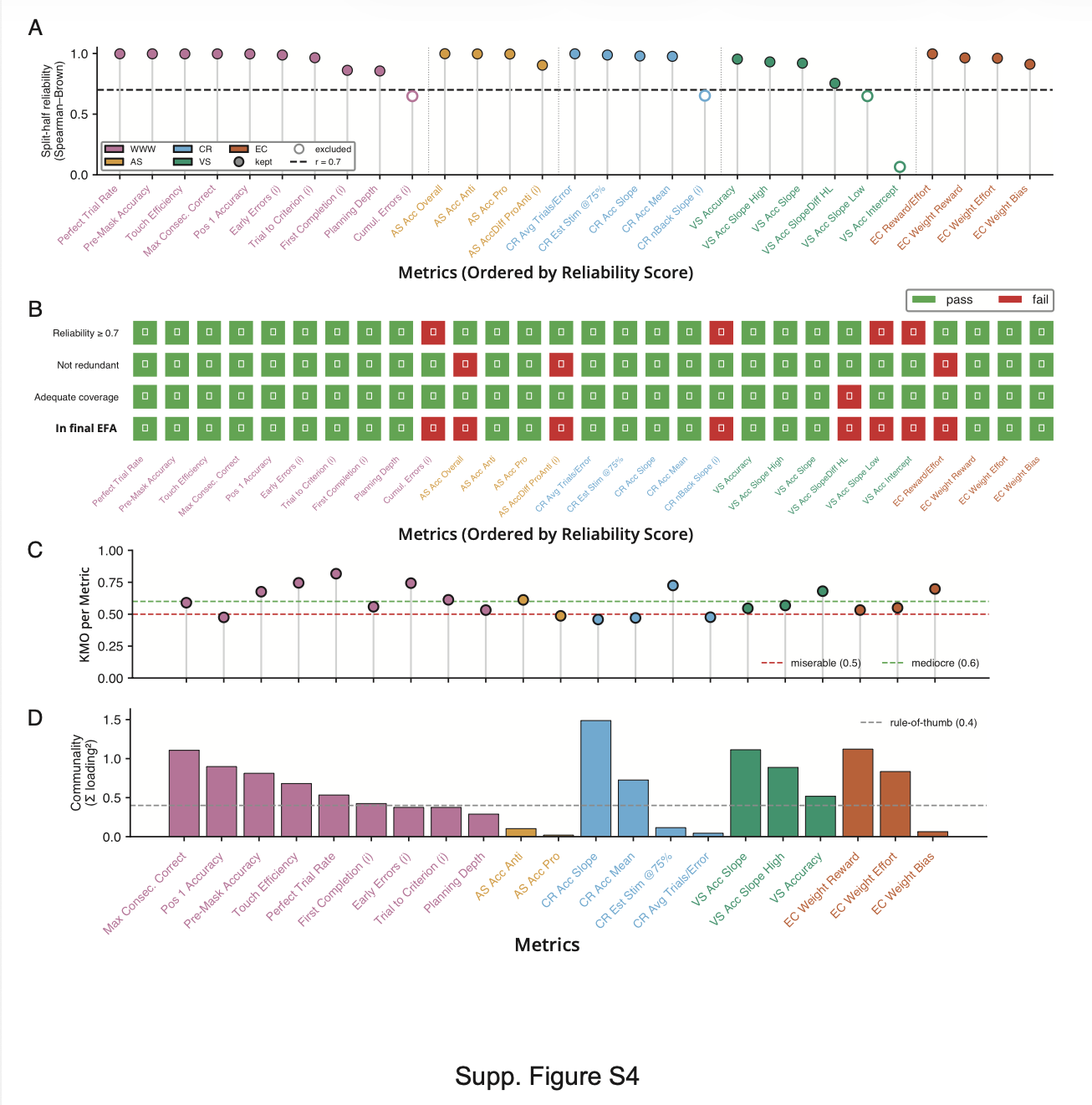

**Supplementary Figure S4. Reliable Metrics Supported Cross-Task Factor Analysis.**

(A) Split-half reliability for all metrics. For each metric, sessions were randomly split into two halves per subject, means computed for each half, and the Pearson correlation between halves across subjects was corrected using the Spearman-Brown formula. This procedure was repeated 100 times and averaged. Dashed line indicates the retention threshold of 0.70. Metrics below this threshold were excluded from subsequent cross-task analyses. (B) Comparison of three screening criteria used to retain metrics for EFA: split-half reliability ≥ 0.70, non-redundancy (pairwise correlation < 0.80; one metric retained from highly correlated pairs), and adequate coverage (metric not missing in most available sessions). Metrics passing all three criteria were included in the cross-task EFA. (C) Per-metric Kaiser-Meyer-Olkin (KMO) measure of sampling adequacy, computed from the within-subject z-scored multitask metric matrix. KMO compares the magnitude of observed correlations to partial correlations for each variable; values closer to 1.0 indicate that correlations are driven by shared latent factors rather than pairwise associations. Variables with KMO < 0.50 are generally considered inadequate for factor analysis. Overall KMO = 0.60. (D) Communality (h²) for each metric from the fitted EFA. Communality is the proportion of a variable's variance explained by the retained factors (h² = 1 − uniqueness). Higher values indicate that the factor solution captures more of that variable's variance. Variables with low communality contribute little to the factor structure.

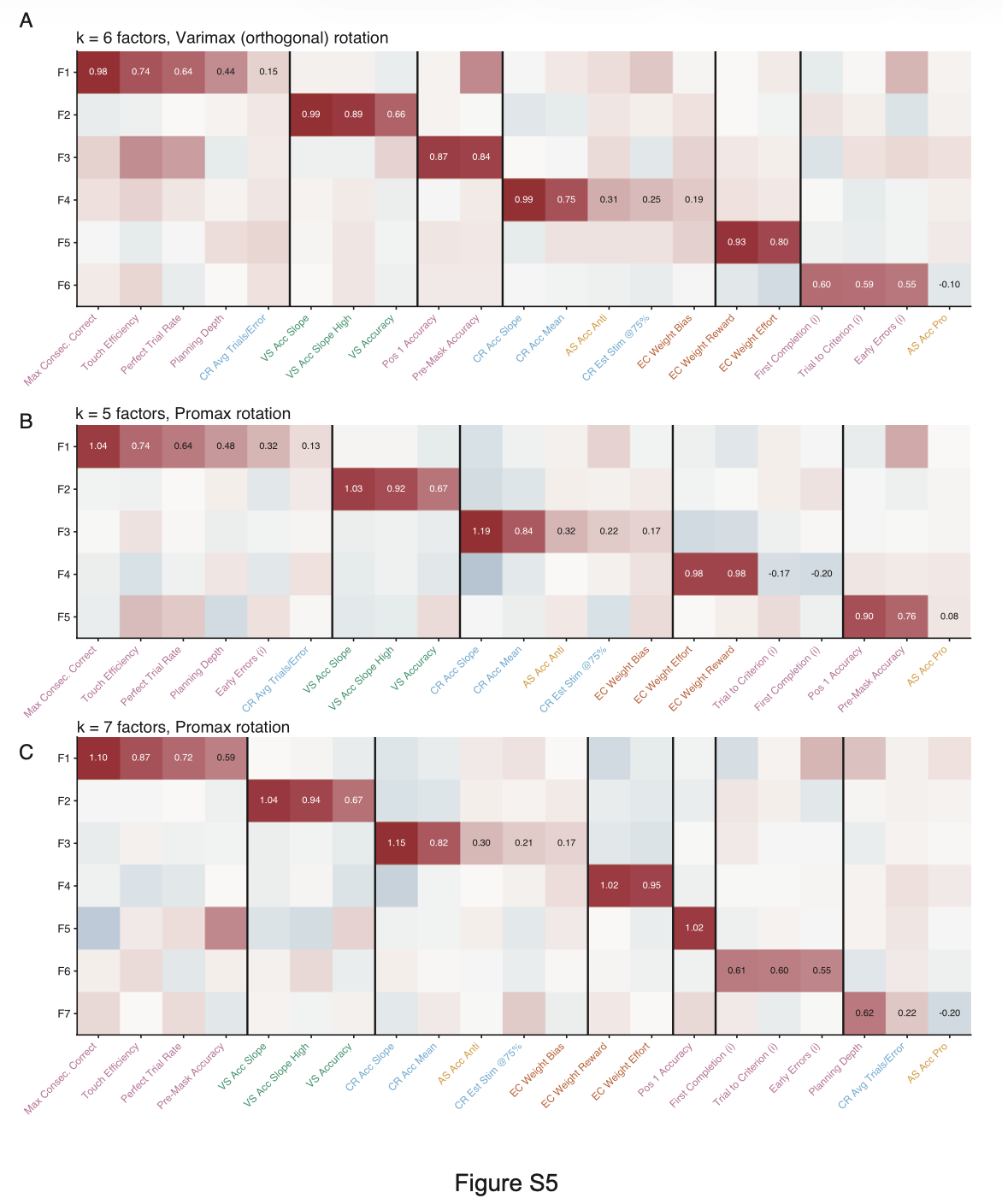

**Supplementary Figure S5. Cross-Task Factor Structure Was Robust Across Analytic Choices.**

(A) EFA with six factors using varimax (orthogonal) rotation, applied to the same pre-processed metric set as Figure 6A. The loading heatmap is compared against the Promax (oblique) solution to assess whether the factor structure is sensitive to rotation method. (B) Sensitivity check with five factors and Promax rotation, applied to the same metric set. Compared against the six-factor solution to assess whether reducing the number of factors merges interpretable components. (C) Sensitivity check with seven factors and Promax rotation, applied to the same metric set. Compared against the six-factor solution to assess whether an additional factor captures meaningful variance or represents noise.

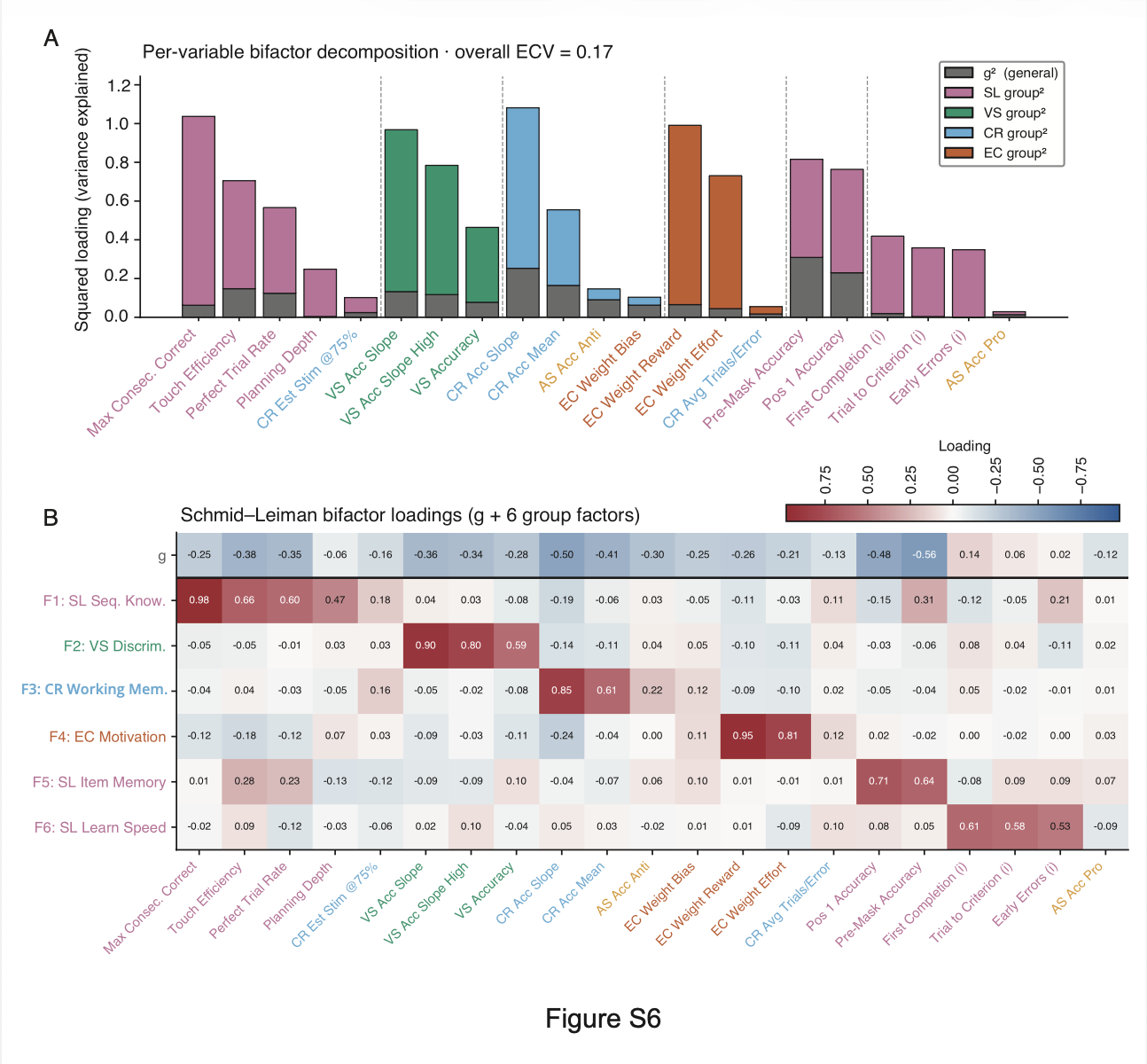

**Supplementary Figure S6. Cross-Task Variance Was Predominantly Domain Specific.**

(A) Per-variable bifactor decomposition via Schmid-Leiman orthogonalization of the Promax solution from Figure 6A. Each variable's explained common variance is partitioned into a general factor (g) component and domain-specific group-factor components, shown as stacked bars (g variance in one color, group-factor variance in another). Per-variable ECV (g / total) is plotted with a 0.50 reference line. Overall ECV = 0.17, indicating that only 17% of the common variance was attributable to the general factor. Most variables showed ECV below 0.50; only EC weight bias and AS accuracy (anti) exceeded 0.50 but remained below 0.70. (B) Schmid-Leiman bifactor loading matrix showing loadings on the general factor (g) plus six orthogonal group factors, displayed in the same variable ordering as Figure 6A. The g loadings were generally weak, while group-factor loadings preserved the task-specific structure seen in the first-order EFA.

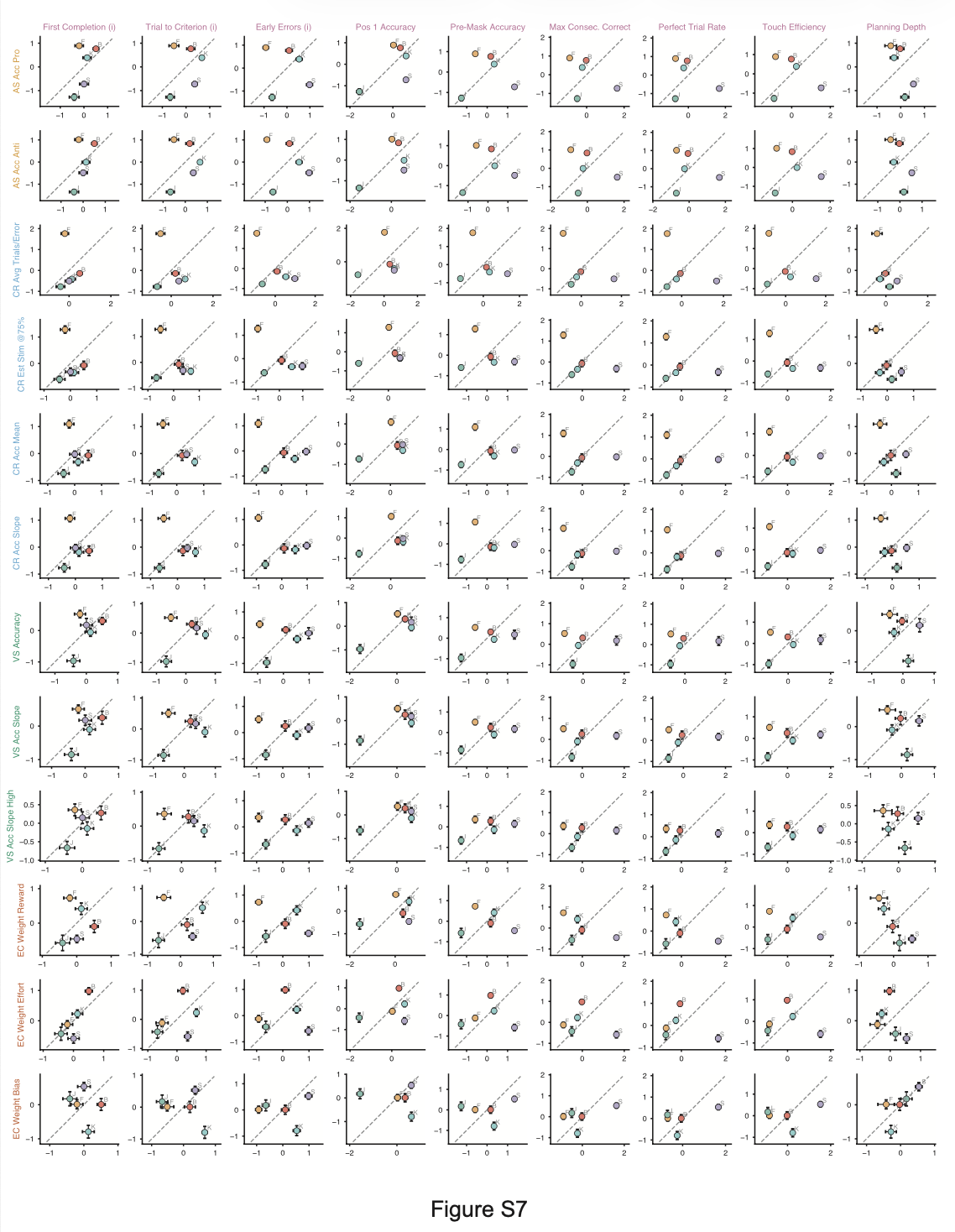

**Supplementary Figure S7. Cross-Task Associations Varied Across Subjects.**

Per-subject session-averaged scatter plots for all sequence learning metrics against all concurrent task metrics, with per-subject Pearson correlations annotated. N = 6 subjects.

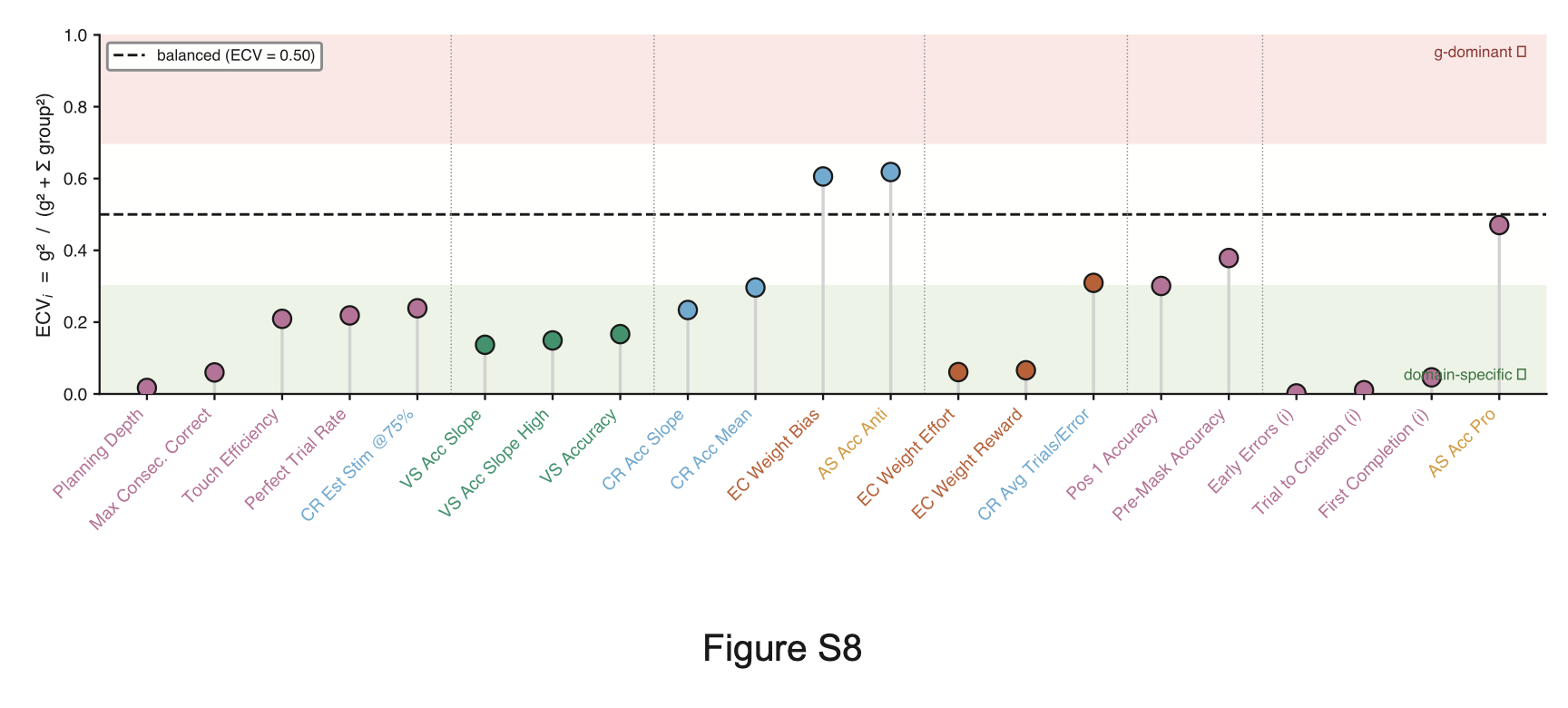

**Supplementary Figure S8. Per-Metric Variance Was Dominated by Domain-Specific Factors.**

(A) Per-variable explained common variance (ECV) ratio (g / total). Most metrics showed ECV below 0.50, indicating that domain-specific factors accounted for more variance than the general factor. Only EC weight bias and AS accuracy (anti) exceeded 0.50 but remained below 0.70, supporting a predominantly domain-specific structure. Overall ECV = 0.17. N = 6 subjects.
